# m6A reader Pho92 is recruited co-transcriptionally and couples translation efficacy to mRNA decay to promote meiotic fitness in yeast

**DOI:** 10.1101/2022.01.20.477035

**Authors:** Radhika A. Varier, Theodora Sideri, Charlotte Capitanchik, Zornitsa Manova, Enrica Calvani, Alice Rossi, Raghu R. Edupuganti, Imke Ensinck, Vincent W.C. Chan, Harshil Patel, Joanna Kirkpatrick, Peter Faull, Ambrosius P. Snijders, Michiel Vermeulen, Markus Ralser, Jernej Ule, Nicholas M. Luscombe, Folkert J. van Werven

## Abstract

*N6-*methyladenosine (m6A) RNA modification impacts mRNA fate primarily via reader proteins, which dictate processes in development, stress, and disease. Yet little is known about m6A function in *Saccharomyces cerevisiae*, which occurs solely during early meiosis. Here we perform a multifaceted analysis of the m6A reader protein Pho92/Mrb1. Cross-linking immunoprecipitation analysis reveals that Pho92 associates with the 3’end of meiotic mRNAs in both an m6A-dependent and independent manner. Within cells, Pho92 transitions from the nucleus to the cytoplasm, and associates with translating ribosomes. In the nucleus Pho92 associates with target loci through its interaction with transcriptional elongator Paf1C. Functionally, we show that Pho92 promotes and links protein synthesis to mRNA decay. As such, the Pho92-mediated m6A-mRNA decay is contingent on active translation and the CCR4-NOT complex. We propose that the m6A reader Pho92 is loaded co-transcriptionally to facilitate protein synthesis and subsequent decay of m6A modified transcripts, and thereby promotes meiosis.

## Introduction

Differentiation from one cell type to another requires accurate and timely control of gene expression. Molecular mechanisms of transcription, RNA processing and translation ensure precise temporal expression of genes. Over the last decade it has become evident that the fate of cells is also largely determined through RNA modifications (Roundtree *et al*, 2017). Specifically, the *N6*-methyladenosine (m6A) modification on messenger RNAs (mRNAs) has been shown to play a pivotal role in cell differentiation, development, and disease pathology (Yang *et al*, 2020).

In mammals, the machinery that deposits m6A, also known as the m6A writer complex, consists of the catalytic subunit METTL3 and the catalytically inactive METTL14. Together they form the RNA binding groove, and are bound by WTAP and several other interacting proteins required to lay down the m6A mark on mRNAs (Liu *et al*, 2014). The m6A writer complex has strong preference for certain motifs (e.g. RRACH in mammals, and RGAC in yeast), and deposits m6A predominantly at the 3’end of transcripts (Dominissini *et al*, 2012; Meyer *et al*, 2012; Schwartz *et al*, 2013). Furthermore, the m6A mark is recognized and bound by reader proteins, which in turn recruit other protein complexes to control the fate of m6A marked transcripts (Zaccara *et al*, 2019). The YT521-B Homology (YTH) domain facilitates the m6A interaction of most m6A reader proteins, which defines the YTH family of proteins that is conserved from yeast to humans including plants (Patil *et al*, 2018). YTH domain containing proteins execute various mRNA fate functions, which includes mRNA decay, translation, transcription, and chromatin regulation (Lasman *et al*, 2020; Wang *et al*, 2014; Xiao *et al*, 2016; Zhou *et al*, 2015). Interestingly, YTH family proteins have been implicated both in translation and decay, suggesting that these proteins may exert overlapping roles in regulating gene expression (Lasman *et al*., 2020; Wang *et al*., 2014; Wang *et al*, 2015; Zaccara & Jaffrey, 2020).

In yeast, the m6A modification occurs during early meiosis, as part of a critical developmental program known as sporulation (Clancy *et al*, 2002; Shah & Clancy, 1992). During sporulation, diploid cells undergo a single round of DNA replication followed by two consecutive nuclear meiotic divisions to produce four haploid spores (Figure 1A). Specifically, the m6A modification occurs during early meiosis (meiotic entry, DNA replication, prophase), and declines once cells undergo meiotic divisions (Schwartz *et al*., 2013) (Figure 1A). The deposition of m6A on mRNAs is catalysed by the methyltransferase Ime4, the Mettl3 orthologue. Ime4 also requires Mum2, the WTAP orthologue, and Slz1, which together comprise the m6A writer machinery in yeast called the MIS complex (Agarwala *et al*, 2012) (Figure 1A). Depending on the strain background, cells lacking Ime4 are either completely or severely impaired in undergoing meiosis and sporulation (Clancy *et al*., 2002; Hongay *et al*, 2006). Some evidence suggests that m6A contributes to the decay and translation of mRNAs (Bodi *et al*, 2015; Bushkin *et al*, 2019). Yeast harbours one known m6A reader protein Pho92, also known as <u>m</u>ethylation <u>R</u>NA <u>b</u>inding protein 1 (Mrb1), which has a conserved YTH domain that is required for its interaction with m6A (Schwartz *et al*., 2013; Xu *et al*, 2015). However, the molecular function of the m6A modification and m6A reader proteins in yeast remains largely unknown.

**Figure 1.**
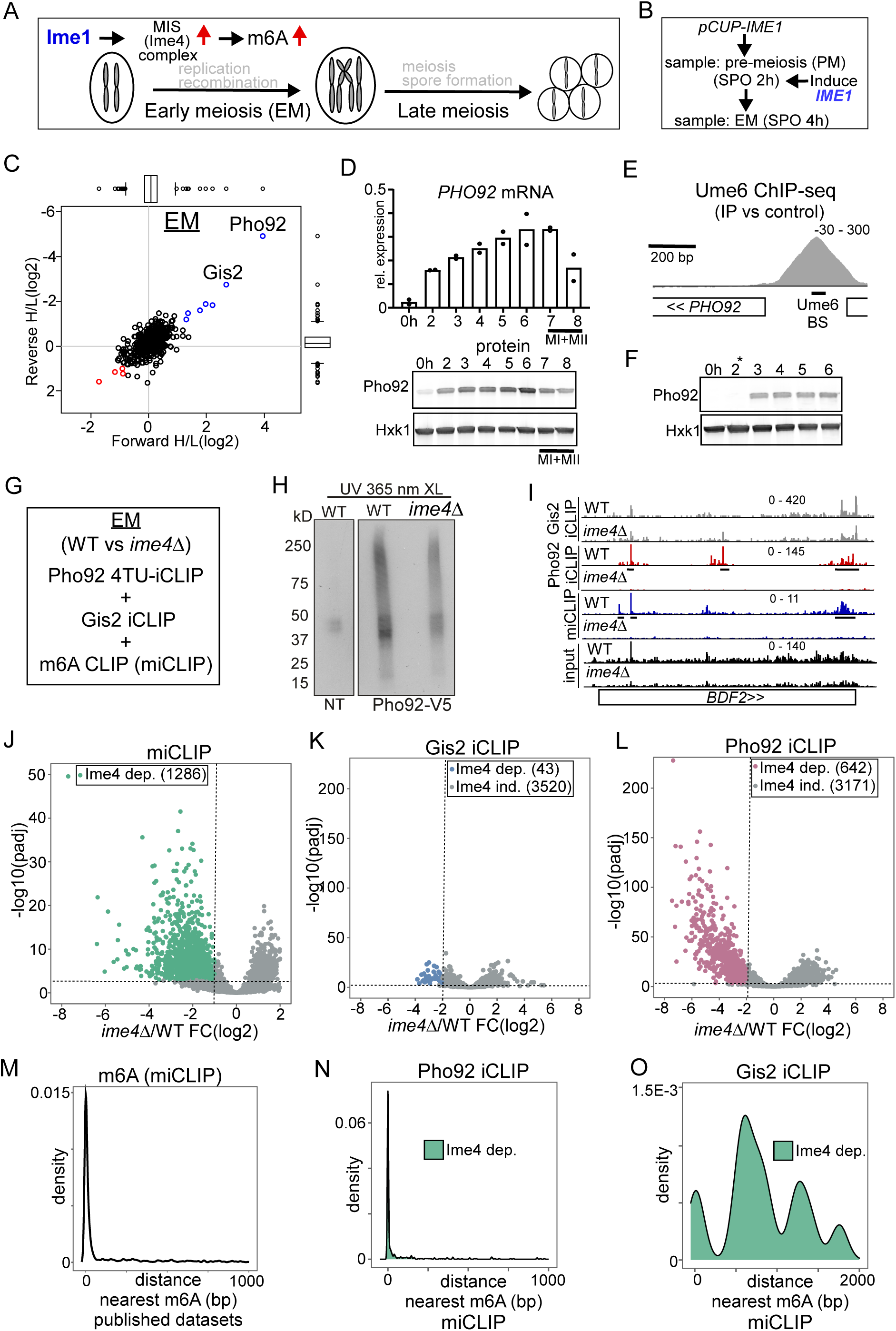
Pho92, but not Gis2, binds to m6A modified transcripts. (**A**) Schematic overview of the yeast meiotic program. Ime1 induces the transcription of the MIS complex. m6A occurs during early meiosis. (**B**) Scheme describing set up for synchronized meiosis. Cells were grown in rich medium till saturation, shifted to SPO, and cells were induced to enter meiosis using *CUP1* promoter fused to *IME1* (*pCUP-IME1*, FW2444). Time points were taken at 2h and 4h. (**C**) Scatter plot displaying proteins identified in m6A consensus oligo pull down versus control. In short, cells were grown in rich medium till saturation, and shifted to SPO, and cells were induced to enter meiosis using *CUP1* promoter fused to *IME1* (*pCUP-IME1*, FW 2444). Protein extracts were incubated using m6A and control RNA oligo bound to streptavidin beads. Eluted proteins were differentially labelled with light and heavy dimethyl isotopes, mixed, and proteins from forward and reverse label swap reactions were identified by MS. (**D**) Pho92 expression prior and during meiosis. Diploid cells with Pho92 tagged with V5 (FW 4478) were induced to enter meiosis in sporulation medium (SPO). Samples were taken at the indicated time points, and Pho92 RNA and protein levels were determined by RT-qPCR and western blotting. (**E**) ChIP-seq data for Ume6 at the *PHO92* locus. Indicated are the Ume6 binding site, and the ChIP-seq signal. Data were taken from *Chia* et al (2021). (**F**) Pho92 expression prior to and after induction of Ime1 expression. Cells harbouring *pCUP-IME1* and Pho92 tagged with V5 (FW 9962) were induced to enter meiosis in sporulation medium (SPO). After 2 hours in SPO, Ime1 was induced with copper sulphate (labelled with *). Samples were taken at the indicated time points, and protein levels were determined by western blotting. Hxk1 was used as loading control respectively. (**G**) Experimental setup for Pho92 4TU-iCLIP, Gis2 iCLIP, and m6A CLIP (miCLIP) in wild-type (WT) and *ime4*Δ cells. (**H**) Autoradiograph showing the protein RNA complexes in no tag (NT) and Pho92-V5 cells. In short, cells were grown till 4 hours in SPO in presence of 4-thiouracil. Cell were harvested and crosslinked. Protein extracts were generated, and Pho92 was immunoprecipitated with anti V5 antibodies. RNA-protein complexes were labelled with (γ-^32^P)-ATP, and separated by SDS page, and transferred to nitrocellulose membrane. Displayed are the signals obtained for no tag control (FW4256), Pho92-V5 (WT, FW 4472), and Pho92-V5 in *ime4*Δ (FW4505). (**I**) Integrative genome browser (IGV) view of *BDF2* gene for Pho92, Gis2 iCLIP and miCLIP in WT and *ime4*Δ cells. Tracks are crosslink per million normalised, strand-specific bigWigs. (**J-L**) Volcano plots comparing WT and *ime4*Δ cells for miCLIP (J), Gis2 iCLIP (K), and Pho92 4TU-iCLIP (L). The Ime4-dependent binding sites are labelled, as determined by a criteria of log2FoldChange <= −1 and adjusted p value < 0.001 for miCLIP and log2FoldChange <= −2 and adjusted *p-*value < 0.001 for the two iCLIP experiments. (**M**) Metanalysis comparing the miCLIP-identified m6A sites to a compendium of published m6A data. For each m6A site the closest published m6A site was calculated. (**N**) Similar as M except that Pho92 binding sites that depended on Ime4 were compared to m6A sites (miCLIP). (**O**) Similar as M except that Gis2 Ime4 dependent sites were compared to m6A sites (miCLIP).

Here we dissected the function of m6A-dependent RNA binding proteins. By employing proteomics and individual-nucleotide resolution UV cross-linking and immunoprecipitation (iCLIP) analysis we reveal that Pho92/Mrb1 is likely the only m6A reader in yeast. Pho92 binds at 3’ ends of mRNAs in both an m6A dependent and independent manner. During early meiosis, Pho92 localizes to the nucleus, through its interaction with the RNA Polymerase II associated transcription elongation complex, Paf1C, and transitions to the cytoplasm to associate with actively translating ribosomes. The Pho92 deletion mutant displays decreased decay of m6A modified mRNAs as well as defects in protein synthesis, indicating a role in translation and decay. We propose that Pho92 is part of an mRNA fate control pathway for m6A modified mRNAs that is instated during transcription and couples translation efficacy to mRNA decay, and thereby promotes gamete fitness in yeast.

## Results

### Pho92/Mrb1 is likely the sole m6A reader in yeast

To systematically identify proteins that interact with m6A in yeast, we incubated m6a modified RNA baits with cell extracts collected either pre-meiosis (PM) or during early meiosis (EM) (Figure 1A and 1B). We used the previously described synchronization method that relies on the induction of the master regulator Ime1 from the *CUP1* promoter (*pCUP-IME1*) to induce meiosis. We collected the PM sample at 2 hours in sporulation medium (SPO) and EM sample at 4 hours in SPO, when the MIS complex is induced by Ime1 and consequently m6A occurs (Figure 1A and 1B) (Agarwala *et al*., 2012; Chia & van Werven, 2016; Schwartz *et al*., 2013). We designed RNA baits consisting of four repeats of the canonical GGACU motif, that were either m6A modified (GGAmCU) or unmodified (GGACU) (Figure S1A) (Edupuganti *et al*, 2017). Subsequently, we performed differential labelling and mass-spectrometry (Figure S1A).

We identified two proteins, Pho92 and Gis2, showing enrichment of binding to m6a modified baits compared to the unmodified bait (log2 enrichment > 2). The known m6A reader and YTH domain containing protein Pho92/Mrb1 was specifically enriched during EM, while Gis2 was enriched in both PM and EM lysates (Figure 1C, Figure S1B, Table S4) (Schwartz *et al*., 2013; Xu *et al*., 2015). Gis2 is an RNA binding protein that associates with ribosomes to facilitate translation by interacting with translation initiation factors (Rojas *et al*, 2012). To validate whether Gis2 preferentially binds to m6A *in vitro* we used recombinant proteins and performed RNA binding assays. Gis2 displayed no binding to either modified or unmodified oligos in a repeating DRACH motif sequence context, though Gis2 binding was detected with a control RNA oligo harbouring its known binding motif of GA(A/U) repeats (Figure S1C and S1D). However, consistent with other reports (Schwartz *et al*., 2013; Xu *et al*., 2015), Pho92 associated with the m6A modified RNA oligo, but not with the unmodified control RNA oligo (Figure S1C). As expected, Pho92 protein lacking the YTH domain (YTHΔ) completely abrogated binding to the m6A modified oligo, whilst removal of the amino-terminus (NΔ), which is largely unstructured, had no impact on m6A modified oligo binding (Figure S1E).

Given that Pho92 was enriched only in early meiosis in the RNA pulldown, we assessed the expression of Pho92 protein and mRNA in wild-type cells entering meiosis. In line with proteomic analysis, we found that Pho92 was expressed in early meiosis prior to when meiotic divisions took place (MI+MII) (Figure 1D). We examined whether the promoter of *PHO92* is possibly under direct control of Ume6, the DNA binding transcription factor that interacts with Ime1 to activate early meiosis gene transcription (Steber & Esposito, 1995). We found that the *PHO92* promoter harbours a canonical Ume6 binding site (CGGCGGCTA) 230 nucleotides (nt) upstream of the start codon, which displays strong binding of Ume6 (Chia *et al*, 2021)) (Figure 1E). To assess whether indeed Pho92 was dependent on Ime1, we induced *IME1* from the *CUP1* promoter (Figure 1B). Prior to *pCUP1-IME1* induction Pho92 was not detectable, but Pho92 transcript and protein rapidly accumulated after *IME1* induction (Figure 1F and S1F). Thus, Pho92 is expressed in early meiosis owing to its promoter being regulated by the meiosis-specific transcription factor Ime1 and its interacting partner Ume6.

### Pho92 associates with mRNAs in an m6A dependent and independent manner

We next set out to determine the transcripts that Pho92 and Gis2 bind *in vivo*, their sequence and positional preferences, and to address whether this interaction is mediated by m6A. Specifically, we used a recently improved iCLIP protocol (Lee *et al*, 2021) to map Pho92 and Gis2 RNA binding sites during early meiosis (4h SPO). Gis2 crosslinked well to RNA with UV irradiation at 254 nm, whereas Pho92 elicited better crosslinking to RNA with UV irradiation at 365 nm in the presence of uracil analog, 4-thiouracil (4TU-iCLIP, Figure 1G, 1H and S1H) (Lee & Ule, 2018). Yeast cells lacking the sole m6A methyltransferase Ime4 display no detectable m6A signal during meiosis (Clancy *et al*., 2002; Schwartz *et al*., 2013). Therefore, we performed iCLIP in both wild-type and *ime4*Δ cells, to distinguish between Ime4-dependent and independent binding. To further assess whether the Ime4-dependent binding equates to specific binding of the reader protein to m6A sites, we performed m6A individual-nucleotide-resolution cross-linking and immunoprecipitation (miCLIP) under the same growth conditions (Lee *et al*., 2021; Linder *et al*, 2015). By using our *ime4*Δ control we were able to distinguish bona fide m6A peaks from non-specific antibody binding. To control for changes in RNA abundance between wild-type and *ime4*Δ conditions, we also produced matched input libraries from mRNAs used for the miCLIP experiments, which also matched the timepoint used for the iCLIP experiments (4h SPO).

Due to the high proportion of cDNAs truncating at the peptide that is crosslinked to RNA fragments, we used the start positions of uniquely mapping reads to assign the positions of crosslink sites in iCLIP, and m6A sites in miCLIP (Lee & Ule, 2018). iCLIP and miCLIP replicate samples were highly reproducible at the level of counts per peak, clustering together in principle component analysis (PCA), with the greatest variance being due to the cells’ genetic background (wild type (WT) or *ime4*Δ) (Figure S1G). However, it is notable that the proportion of variance accounted for by genetic background was much less for Gis2 (34%) than for the miCLIP (80%) or Pho92 (58%). For example, the *BDF2* transcript harboured several Pho92 binding sites that depended on Ime4, while Gis2 displayed m6A independent binding (Figure 1I).

We found 1286 miCLIP peaks in 870 genes that were reduced in *ime4*Δ cells (log2FoldChange <= −1, adjusted p value < 0.001), which we will hereby refer to as m6A sites (Figure 1J). Analysis of Gis2 iCLIP revealed 3563 peaks, of which only 43 peaks in 24 genes were decreased in *ime4*Δ representing only 1.2% of all Gis2 peaks (log2FoldChange <= −2, adjusted p value < 0.001) (Figure 1K). A much larger number of Pho92 peaks (642 peaks in 507 genes, 16.7% of all detected peaks) were found to be reliably reduced in *ime4*Δ (log2FoldChange <= −2, adjusted p value < 0.001) (Figure 1L). Using a less stringent criteria we could designate up to 30% (1130/3823 peaks at log2FoldChange <= 0, adjusted p value < 0.05) of detected Pho92 peaks as reduced in *ime4*Δ, however for subsequent analysis we will refer to the 642 stringently defined peaks as “Ime4-dependent” Pho92 binding sites. This means that surprisingly, a large subset of Pho92 binding sites do not decrease in *ime4*Δ cells, indicating that Pho92 can also associate with transcripts in an Ime4 independent manner (Figure 1L). We ascertained that this is likely not background binding of Pho92 because in *ime4*Δ cells clear enrichment for Pho92-RNA complexes can be detected compared to the untagged control (Figure 1H and S1H). Indeed, when we implemented the iCLIP protocol to the untagged control, we found overall much less signal compared to the Pho92 iCLIP (Figure S1I).

To further explore whether Ime4-dependent Gis2 and Pho92 binding was also m6A-dependent, we calculated the distance to the nearest m6A site from each Ime4-dependent Gis2 and Pho92 binding site. To do so, we first validated our miCLIP data against published datasets containing analyses of multiple time points during meiosis using different m6A-sequencing techniques (Garcia-Campos *et al*, 2019; Schwartz *et al*., 2013). Even though the time point of our miCLIP and the published datasets were not a match, m6A sites defined by miCLIP overlapped with m6A sites from the published datasets with ∼30% of miCLIP m6A sites being within 50nt of a published site (Figure 1M and S1J). Similarly, Ime4-dependent Pho92 sites mapped close to m6A sites as determined by miCLIP or published data (Figure 1N and S1K). However, the few Ime4-dependent Gis2 binding events showed no clear proximity to m6A sites defined by either methodologies (Figure 1O and S1L). Further scrutiny of Pho92 binding revealed that the Ime4 dependent Pho92 association signal is highly enriched at the m6A sites we identified using miCLIP (Figure S1M). We conclude that a substantial portion of Pho92 binding is dependent on m6A, while Gis2 binding to RNAs does not require the presence of m6A. Taken together, our data strongly suggest that Pho92 is likely the only m6A reader protein in yeast. For the remainder of this study we focus on our efforts to reveal the mechanism by which Pho92 controls the fate of m6A marked transcripts during yeast meiosis.

### Features of transcripts bound by Pho92

Upon closer inspection of the Pho92 iCLIP data, we found that Pho92 iCLIP peaks are found in the mRNAs of key regulators of early meiosis such as *IME1* and *RIM4* (Figure 2A). The *IME1* and *RIM4* Pho92 iCLIP peaks overlapped with m6A sites identified with miCLIP (*RIM4*), or, in the case of *IME1*, with comparative Nanopore sequencing of WT versus *ime4*Δ described previously (Leger *et al*, 2021) (Figure S2A). As an exception, *IME2,* a key kinase in meiosis, displayed no detectable Pho92 binding, but, consistent with previous reports, it did however display an Ime4-dependent peak in miCLIP data, albeit below our stringent thresholds (log2FC=-0.6, padj= 0.007) (Bodi *et al*, 2010; Schwartz *et al*., 2013) (Figure 2B). There was a good overlap (more than 50%) between Pho92 binding sites and m6A sites (Figure 2C). As noted for *IME2*, there were also m6A sites where Pho92 was not detected, which is likely technical in nature as our current data coverage is not sufficient to cover all m6A sites with either miCLIP or Pho92 iCLIP, and additional variation could be caused due to technical differences between miCLIP and iCLIP protocols.

**Figure 2.**
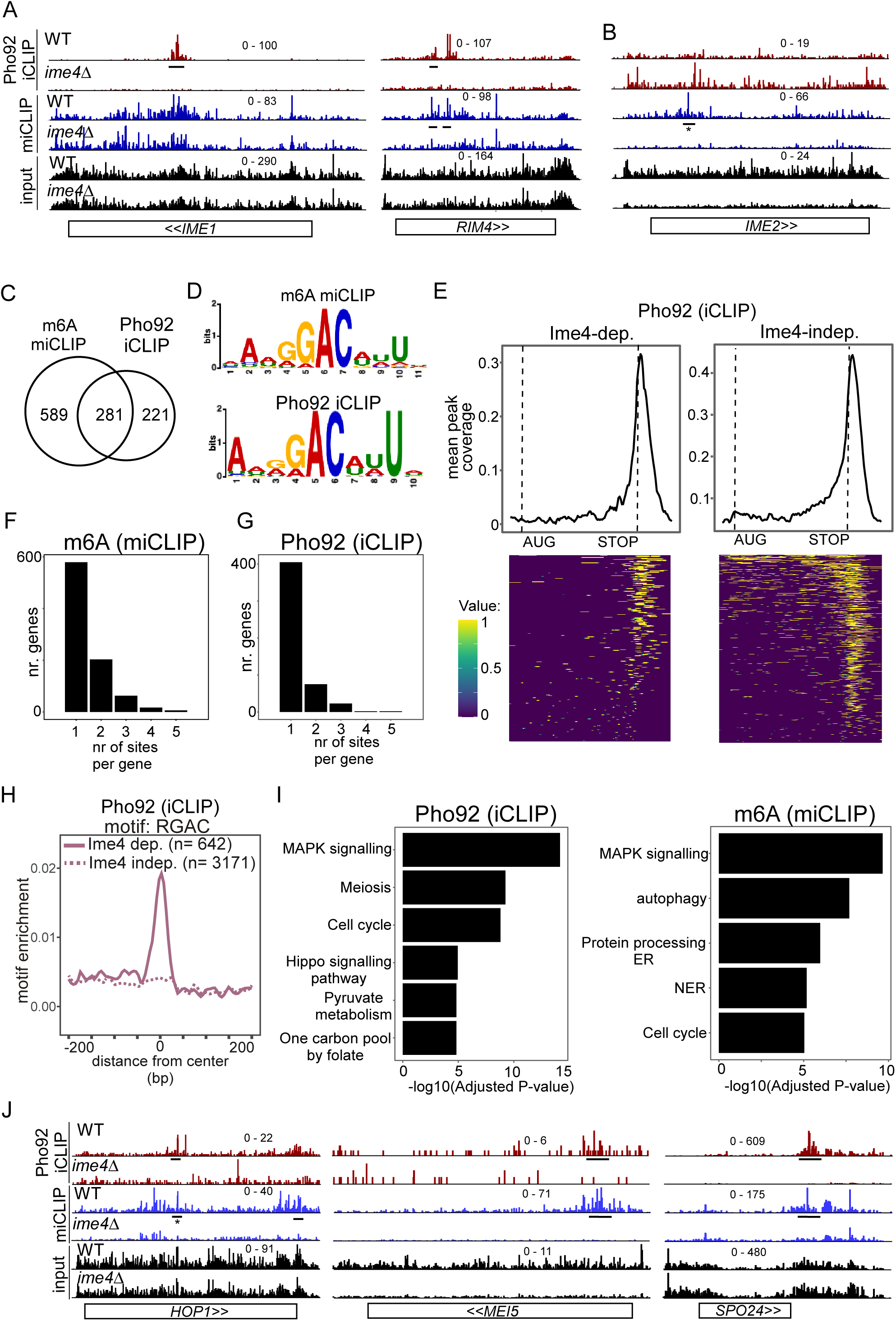
Features of transcripts bound by Pho92. (**A**) *IME1* and *RIM4* transcripts bound by Ime4-dependent Pho92. Shown are the crosslinks per million normalised bigWigs for Pho92 iCLIP and miCLIP in WT and *ime4*Δ cells. Underlined are the binding sites identified in the analysis. (**B**) Similar as A, except that *IME2* locus is displayed, which exhibits an m6A peak but no Pho92 binding. The m6A site is labelled. * this was reported in our analysis below the significance threshold (log2FC=-0.6, padj= 0.007). (**C**) Venn diagram showing the overlap between genes with m6A sites and Ime4-dependent Pho92 binding. (**D**) Sequence logos of the top ranked motifs from m6A sites and Ime4-dependent Pho92 sites as determined by STREME. (**E**) Pho92 metagene profile split into genes containing Ime4-dependent Pho92 binding sites (left) and Ime4-independent sites (right). The matrix underlying the heatmap is scored 1 for binding site and 0 for no binding, intermediate values between 0 and 1 are due to smoothing in the visualisation. (**F**) Number of m6A sites (miCLIP) per transcript. On the x-axis transcripts with 1, 2, 3, 4 or 5 bindings sites. On the y-axis the number of genes for each category is displayed. (**G**) Similar to F, except that Pho92 binding sites were analysed. (**H**) Motif enrichment around m6A sites and Pho92 Ime4-dependent and independent binding sites for RGAC. Motif frequency plotted around the centre of Pho92 binding sites, split into Ime4-dependent sites (solid line) and Ime4-independent sites (dotted line). (**I**) Gene ontology (GO) enrichment analysis for Pho92 bound transcripts (left) and m6A harbouring transcripts based on the miCLIP analysis (right). On the y-axis the category of processes involved, while on x-axis the -log10(adjusted *P*-value) is displayed. (**J**) *HOP1, MEI5, and SPO24* transcripts bound by Pho92 and marked with m6A. Data tracks are crosslinks per million normalised bigWigs, visualised in IGV. Underlined are the binding sites identified in the analysis. * this was reported in our analysis below the significance threshold (log2FC=-0.94, padj= 0.004).

Using STREME, we analysed the RNA sequence context of Ime4-dependent Pho92 binding. Strikingly, the highest ranked motif found in Ime4-dependent Pho92 binding sites was nearly identical to the highest ranked motif found for the m6A sites, which highlights again that the Ime4-dependent sites are mostly m6A-dependent sites (Figure 2D, S2B and S2C). This motif is an “extended” RGAC sequence context, containing upstream AN and downstream NNU nucleotides that were previously described in (Schwartz *et al*., 2013). The Ime4-independent binding sites of Pho92 showed no enrichment for the m6A motif sequence but showed weak enrichment for short motif sequences containing GU dinucleotides, indicating that m6A independent binding of Pho92 is conceivably driven independent of sequence context (Figure 1H, S2B and S2C).

As previously stated, consistent with the m6A pattern at mRNAs, Pho92 binding was detected predominantly at the 3’ end of transcripts, with 23% (145/642) of Ime4-dependent binding sites and 13% (410/3171) of Ime4-independent binding sites directly overlapping a STOP codon (Figure 2E and S2D). We also found Pho92 binding sites in the protein coding sequence or at the 5’ end of transcripts, which included key meiotic mRNAs (*IME1* and *RIM4*) (Figure 2A and S2D). Although most transcripts contained just one m6A and/or a Pho92 binding site, some transcripts did contain two or more m6A and/or Pho92 binding sites and that 3’ end binding of Pho92 is at least partially independent of m6A and the m6A consensus sequence motif (Figure 2F and 2G). The Ime4-independent binding sites also showed positional enrichment at the 3’ end of mRNA transcripts, suggesting that these are likely bona fide binding sites (Figure 2E and S2D). Thus, Pho92 associates with transcripts predominantly at their 3’ ends, and a subset of this binding occurs in an Ime4-dependent manner at m6A sites within canonical m6A motifs.

We performed gene ontology (GO) analysis of Pho92 bound transcripts (Ime4-dependent) and found that MAPK signalling, cell cycle and meiosis were the top enriched terms (Figure 2I). Notably, examination of m6A sites showed enrichment for MAPK signalling and cell cycle but not meiosis (Figure 2I), suggesting that m6A by itself regulates a different subset of transcripts than Pho92. Genes that showed Ime4-dependent Pho92 binding and corresponding m6A sites included those involved in regulating the meiotic program (such as *IME1*, *RIM4* and *SPO24),* DNA recombination and double strand-break repair (such as *HOP1* and *MEI5*) (Figure 2J, 2A and S2A).

### Pho92 is important for the onset of meiosis and fitness of gametes

The yeast meiotic program is regulated by the master regulatory transcription factors, Ime1 and Ndt80 (Chu *et al*, 1998; Kassir *et al*, 1988). Ime1 drives early meiosis, while Ndt80 regulates the genes important for late meiosis, which includes meiotic divisions and spore formation (Figure 1A) (van Werven & Amon, 2011). GO-analysis of Pho92 bound transcripts revealed that Pho92 associates with genes important for meiosis. This posed the question whether Pho92 acts at the same genes as Ime1 and Ndt80, or perhaps regulates a different category of genes, independent of these two transcription factors. To identify Ime1 and Ndt80 regulated genes we performed RNA-seq in *ime1*Δ and *ndt80*Δ cells in early meiosis (4 hours in SPO). As expected, the genes downregulated in *ndt80*Δ showed strong enrichment for the Ndt80 motif sequence, indicating the RNA-seq was able to identify direct target genes (Figure S3A, S3B)(de Boer & Hughes, 2012). The promoter of transcripts downregulated in *ime1*Δ, however, showed enrichment for both Ndt80 and Ume6 motifs, likely because *ime1*Δ blocks induction of genes indirectly regulated by Ime1 (Figure S3A). We compared the genes that were significantly downregulated in *ime1*Δ and *ndt80*Δ cells to Pho92 bound transcripts. While the promoters of Pho92 bound transcripts (iCLIP) were enriched for the Ume6 motif, we found that only 81 genes out of the 507 Pho92 targets overlapped with the 1133 Ime1 regulated transcripts (Figure 3A and S3A). Moreover, 54 genes overlapped with the 695 Ndt80 regulated transcripts (Figure 3A). Thus, a large fraction of transcripts bound by Pho92 were likely not targets of Ime1 or Ndt80. Similarly, little overlap was observed between m6A-marked transcripts and transcripts regulated by Ime1 or Ndt80 (Figure S3C). Our data suggest that m6A and Pho92 regulate a distinct subset of transcripts or genes than the transcription factors (Ime1 and Ndt80) that control the meiotic program.

**Figure 3.**
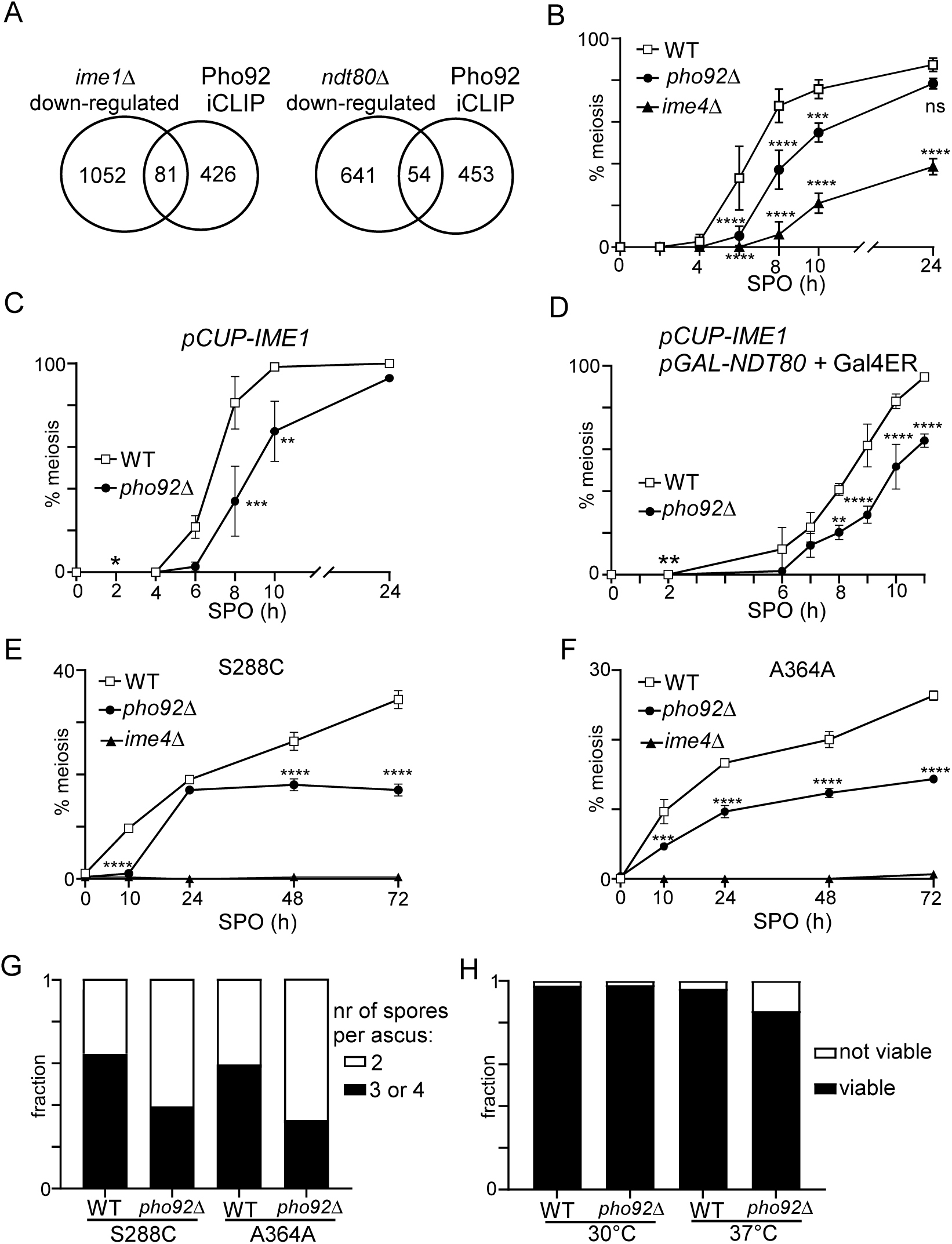
Pho92 is important for meiosis and fitness of gametes. (**A**) Venn diagram displaying the comparison between RNA-seq of *ime1*Δ (FW81) and *ndt80*Δ (FW 4911), and transcripts bound by Pho92. For this analysis, Ime1 and Ndt80-dependent genes were selected by taking the transcripts that were significantly down-regulated compared to the WT control. (**B**) Onset of meiosis in WT, *pho92*Δ, and *ime4*Δ cells (FW1511, FW3528, and FW725). Cells were grown in rich medium till saturation, shifted to pre-sporulation medium and grown for an additional 16 hours. Subsequently, cells were shifted to SPO, and samples were taken at the indicated time points. Cells were fixed, and stained, DAPI masses were counted for at least n=200 cells per biological repeat. Cells with two or more DAPI masses were considered as undergoing meiosis. (**C**) Similar as B, except that cells were induced to undergo meiosis using *pCUP-IME1* (WT FW2444, *pho92*Δ FW3576). Cells were grown as described in B, shifted to SPO, and after 2 hours treated with copper sulphate to induce *IME1* expression. (**D**) Similar as C, except that these also harboured *NDT80* under control of the *GAL1-10* promoter and fused Gal4 activation domain plus estrogen receptor (Gal4ER). Ndt80 expression was induced at 2 hours in SPO with β-estradiol together with Ime1 expression (WT FW2795, *pho92*Δ FW9070). (**E**) Similar as B, except that S288C strain background was used for the analysis (WT FW631, *pho92*Δ FW8983, *ime4*Δ FW8985). (**F**) Similar analysis as B, except that A364A was used for the analysis (WT FW1671, *pho92*Δ FW8912, *ime4*Δ FW8913). The error bars represent the standard error of the mean (SEM) of n = 3; *p< 0.05 **p< 0.01 ***p < 0.001, ****p < 0.0001, compared to WT control on a two-way ANOVA followed by a Fisher’s least significant difference (LSD) test. (**G**) Spore packaging was assessed in WT and *pho92*Δ strains (S288C and A364A strains as in E and F). Number of packaged spores per ascus were counted for at least 200 asci. (**H**) Spore viability of WT (FW 1511) and *pho92*Δ (FW3531). Cells were patched from YPD agar plates to SPO agar plates and incubated for 3 days at 30°C or 37°C. Subsequently tetrads were dissected, and spores grown on YPD agar plates. The fraction of viable and not viable spores are indicated. At least n=150 spores for each condition (30°C or 37°C) and each strain (WT and *pho92*Δ) were used for the analysis.

Given that Pho92 is specifically expressed during early meiosis (Figure 1D), we expect that, like Ime4, Pho92 plays a role in controlling gametogenesis. Therefore, we closely examined the effects of *pho92*Δ on meiosis and spore formation in the SK1 strain background, widely used for studying meiosis and sporulation. We found that *pho92*Δ exhibits a delay in the onset of meiosis compared to WT, though much milder than *ime4*Δ (Figure 3B). This is likely due to non-catalytic functions of Ime4, suggested by a milder phenotype of Ime4 catalytic mutant vs WT protein {Agarwala, 2012 #11}. We found that the delay in meiotic divisions observed in *pho92*Δ cells was not alleviated by inducing *pCUP-IME1* (Figure 3C), supporting the idea that Pho92 and Ime1 regulate different subsets of genes. We also assessed whether Ndt80 expression can alleviate the effect of *pho92*Δ on meiosis. To do so, we expressed Ime1 and Ndt80 together when cells were starved in sporulation medium (SPO) (Figure 3D). We induced Ndt80 from the *GAL1-10* promoter (*pGAL-NTD80*) using the transcription factor Gal4 fused to estrogen receptor (Gal4ER) and addition of β-estradiol (Benjamin *et al*, 2003; Chia & van Werven, 2016). Expressing Ime1 and Ndt80 together surprisingly had little effect on meiosis, as most cells (more than 80%) completed at least one meiotic division by 8 hours post-induction (WT, Figure 3D). Approximately 50% of *pho92*Δ cells completed one meiotic division after 8 hours of Ime1/Ndt80 induction (Figure 3D). Thus, the delay in meiosis in *pho92*Δ cells was not dependent on the expression of Ime1 and Ndt80.

We further assessed whether Pho92 contributes to the onset of meiosis in strain backgrounds that sporulate with a lower efficacy (S288C and A364A). Both S288C and A364A displayed more than 50 percent less meiotic cells after 72 hours in SPO (Figure 3E and 3F). We further found that Pho92 contributes to the packaging of spores because the fraction of tetrads per asci decreased and the fraction of dyads increased in *pho92*Δ cells (Figure 3G). Finally, we quantified the effect of Pho92 on spore viability itself. We observed a modest reduction of spore viability when cells underwent sporulation at elevated temperatures (37°C) (Figure 3H). We conclude that the m6A reader Pho92 is important for the fitness of meiotic progeny in yeast.

### Paf1C retains Pho92 in the nucleus during early meiosis

YTH domain containing proteins are conserved from humans to plants, and exercise various functions to control the fate of m6A transcripts (Patil *et al*., 2018). Little is known about the role of Pho92 in regulating the fate of m6A modified transcripts. To identify a possible function for Pho92 in yeast, we determined its protein-protein interactions. We performed immunoprecipitation of Pho92 followed by label-free quantitative mass spectrometry. In addition to purifying Pho92 from cells staged in early meiosis, we also expressed Pho92 from heterologous promoters to intermediate levels (*CYC1* promoter, *pCYC1*-FLAG-Pho92) in cycling cells (Figure 4A, S4A-S4B). The purification of Pho92 from cells staged in early meiosis identified few interacting proteins and weak enrichment for Pho92 itself, likely because of the high protein turnover in extracts of meiotic cells (Figure S4A and S4C). In contrast, the purification from cycling cells grown in YPD identified large set of proteins that co-purified with Pho92 (Figure S4B). Specifically, several proteins involved in ribosomal biogenesis and translation were enriched compared to the untagged control (Figure 4A and Table S6). Surprisingly, we also found that Pho92 interacted with proteins involved in chromatin organization and transcription. These included two subunits of RNA Polymerase II (pol II) associated transcription elongation complex Paf1C (Leo1 and Paf1), as well as the highly conserved and essential pol II associated transcription elongation factor Spt5 (Figure 4A).

**Figure 4.**
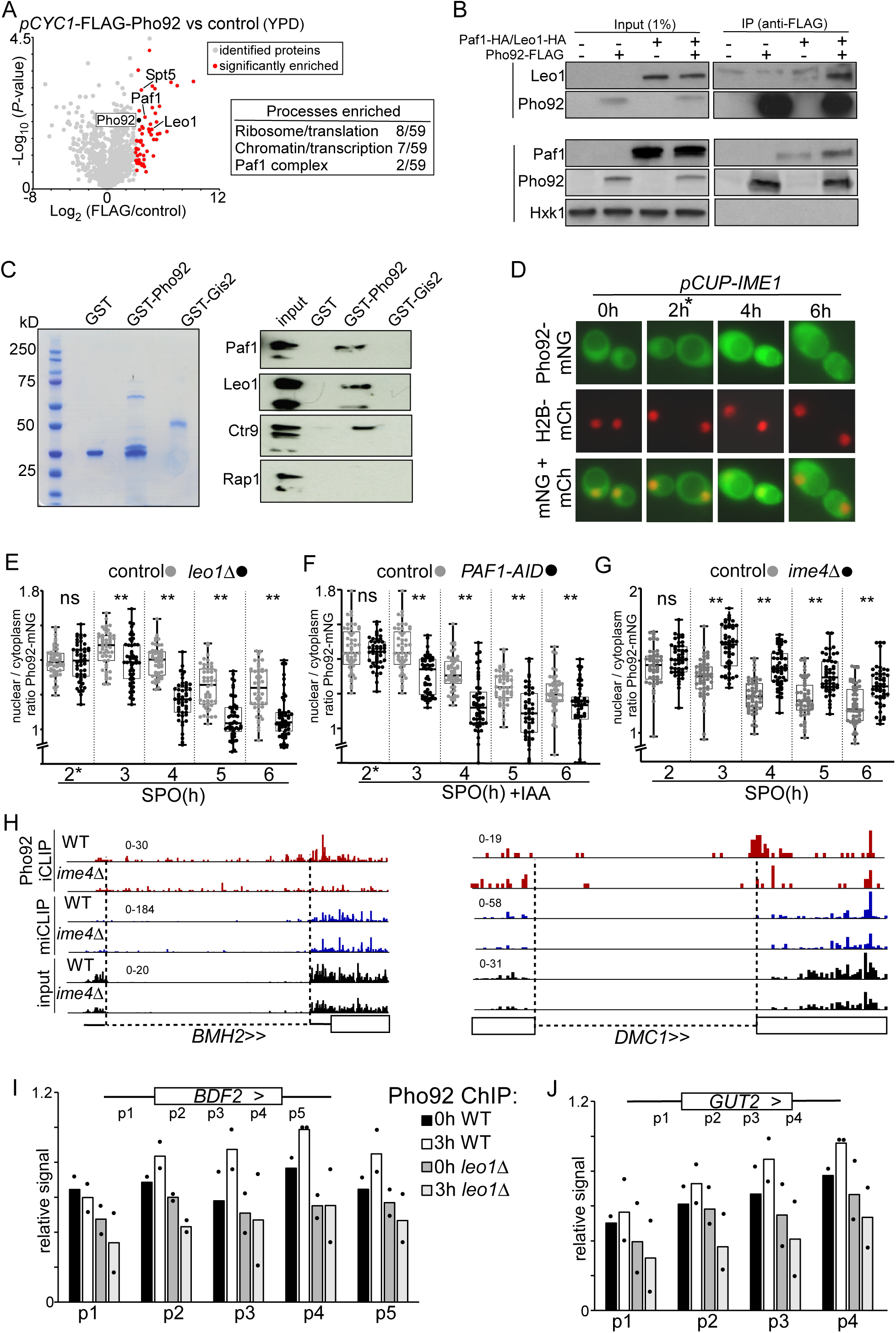
Paf1C interacts with Pho92 to direct Pho92 to nucleus. (**A**) Volcano plot IP-MS of Pho92 (left). Cells expressing the *CYC1* promoter (*pCYC1-*FLAG-Pho92, FW8734) controlling Pho92 expression and a FLAG-tag at the amino terminus were grown in rich medium conditions. Cell extracts from *pCYC1*-FLAG-Pho92 and control (FW629) were incubated with anti-FLAG beads and eluted with FLAG peptides and analysed by MS using label free quantification method. Significantly enriched proteins are displayed in red. Table (right) proteins enriched in IP-MS experiment (also data from plots in Figure S4). Highlighted are the processes and protein complexes enriched. (**B**) Co-immunoprecipitation of Pho92 and Paf1C. We used strains where *pPYK1*-FLAG-Pho92 was expressed with or without tagged HA-Paf1 or HA-Leo1 (FW9880 and FW9791). Cells harbouring FLAG-Pho92, HA-Paf1, or HA-Leo1 only were used as control (FW8732, FW9782 and FW9763). As a negative, control membranes were also probed for Hxk1, which does not interact with Pho92. (**C**) Pull-down of Paf1C by GST-Pho92, GST-Gis2 or GST alone induced in bacteria. Cell extracts were prepared and immobilized on Glutathione-agarose beads. GST-Pho92 bound to beads were subsequently incubated with extracts expressing HA-tagged Paf1, Leo1, Crt9, and Rap1 (FW9782, FW9763, FW9784, FW 4948). (**D**) Localization of Pho92 during entry in meiosis. Cells expressing Pho92 fused with mNeongreen (Pho92-mNG, FW9633) were used for the analysis. To determine nuclear localization, we used histone H2B fused to mCherry (H2B-mCh) and the cells also harboured *pCUP-IME1* to enable induction of synchronous meiosis. Cells were treated with copper sulphate at 2 hours (labelled with *), and samples were taken at the indicated time points. (**E**) Quantification of nuclear over cytoplasmic signal for Pho92-mNG in WT and *leo1*Δ (FW9633 and FW9736). Cells were grown as described in D. At least 150 cells were quantified per time point. Each datapoint is shown in addition to the box plots. ** *p* < 0.01 Welch’s paired t test. (**F**) Similar to E, except with Paf1 depletion strain. Paf1 is fused to an auxin induced degron (AID-tag), which is induced by copper sulphate and IAA treatment at 2h (FW10128). These cells also express *TIR1* ligase under control of the *CUP1* promoter. Cells harbouring the *TIR1* ligase alone were used as controls (FW10129). (**G**) Same as in E, except that *ime4*Δ cells were used for the analysis (FW9604). (**H**) Pho92 binding overlaps with *BMH2* and *DMC1* introns. Data tracks of Pho92 iCLIP, miCLIP, and miCLIP-input in WT and *ime4*Δ cells. Intron regions are designated with dashed lines. (**I**) ChIP-qPCR of Pho92 at the *BDF2* and *GUT2* loci during entry into meiosis in WT (FW4478) and *leo1*Δ (FW10113) cells. Biological repeats of WT and *leo1*Δ cells were grown in parallel, each sample was input normalized, subsequently the primer pair with highest signal was set to 1 for each biological repeat, which was primer pair p4 for 3h WT *BDF2* and *GUT2*. The relative mean signal of n=2 biological repeats are displayed.

To validate the interaction between Pho92 and Paf1C, we co-expressed epitope-tagged Pho92 and Paf1C and performed co-immunoprecipitation of Pho92 with Paf1C subunits Leo1 and Paf1 followed by western blotting. We found that Leo1, and to a lesser extent Paf1, co-immunoprecipitated with Pho92, but not the negative control hexokinase 1 protein (Figure 4B). Additionally, we affinity purified Pho92 fused to GST from *E. coli* and determined whether yeast extracts with HA-tagged Paf1, Leo1 or Ctr9 associated with recombinant GST-Pho92. We found that Paf1C subunits (Paf1, Leo1 and Ctr9) were enriched in the GST-Pho92 pull-down compared to the GST only or GST-Gis2 negative control, while the transcription factor Rap1 did not show enrichment (Figure 4C). We conclude that Pho92 interacts with Paf1C *in vitro* and *in vivo*.

Paf1C associates with pol II to promote transcription elongation and chromatin organization during transcription (Rondon *et al*, 2004; Van Oss *et al*, 2017). If Pho92 and Paf1C interact during meiosis, we hypothesized that Pho92 must localize to the nucleus and possibly associate with gene bodies during transcription. To address this, we first determined whether the Paf1C regulates Pho92 localization in cells entering meiosis. To visualize Pho92 we tagged the protein with mNeongreen (Pho92-mNG) and monitored Pho92-mNG localization during a synchronized meiosis (*pCUP-IME1*). We determined nuclear Pho92 signal using histone H2B fused to mCherry (H2B-mCh) (Figure 4D). A large fraction of Pho92-mNG signal was in the nucleus prior to induction of meiosis (0h and 2h) and in the early time points of meiosis (3 and 4 hours) (Figure 4D and 4E (see WT)). At the later time points (5 and 6 hours), Pho92-mNG was however, largely excluded from the nucleus (Figure 4D and 4E (see control)). It is noteworthy that the Pho92-mNG whole cell signal was lower prior to induction of meiosis (Figure S4D-S4F). We conclude that Pho92 localization is dynamically regulated in cells undergoing early meiosis.

Next, we examined whether Paf1C contributed to Pho92 localization to the nucleus. We visualized Pho92-mNG localization in *leo1*Δ cells and in Paf1 depleted cells using the auxin induced degron (*PAF1-AID*) (Figure S4G). From 3 hours into sporulation onwards, we found that Pho92 was less nuclear in *leo1*Δ cells compared to the control (Figure 4E). A comparable difference in nuclear localization of Pho92 was observed when we depleted Paf1 (*PAF1-AID* + IAA) (Figure 4F). We also examined whether Pho92 localization was dependent on the m6A writer Ime4. In *ime4*Δ cells Pho92 was retained more in the nucleus, indicating that m6A modified transcripts normally facilitate its exit from the nucleus (Figure 4G). We conclude that Pho92 localization is regulated throughout early meiosis. While Paf1C contributes to localization of Pho92 to the nucleus, m6A modified transcripts facilitate transition of their reader Pho92 into the cytoplasm.

### Paf1C directs Pho92 to target genes

Interaction between Leo1 and nascent mRNA has been reported to stabilize the association of Paf1C with transcribed genes (Dermody & Buratowski, 2010). Our data showing the effect of Paf1C on Pho92 localization to the nucleus raised the question of whether Pho92 is recruited to pre-mRNAs during transcription via Paf1C. To corroborate this idea that Pho92 associates with nascent mRNAs we scanned the iCLIP data for Pho92 association with intron regions which are typically retained in nascent mRNAs. We found 55 transcripts with Pho92 iCLIP signal in the intronic regions. For example, *BMH2* and *DMC1* intron regions contained Pho92 iCLIP crosslinks, while miCLIP and input displayed no detectable crosslinking within the intron sequence (Figure 4H and S4H).

To substantiate the observation that Pho92 associates with some of its targets during transcription, we performed Pho92 chromatin immunoprecipitation (ChIP) qPCR across two genes that contained Ime4-dependent Pho92 binding sites (*BDF2* and *GUT2*) (Figure 1I and 6J). In wild-type cells prior to entering meiosis (0 hours SPO), we found some Pho92 enrichment at these two ORFs (Figure 4I and 4J). Cells entering meiosis (3h SPO) showed an increase in Pho92 binding. Importantly, in *leo1*Δ cells Pho92 ChIP signal over the whole gene body was reduced (Figure 4I and 4J). This decrease cannot be attributed to Pho92 expression differences or meiotic defects because *leo1*Δ cells entered meiosis with a mild delay and expressed Pho92 to comparable levels as wild-type cells (Figure S4I and S4J). Also, the Paf1C deletion or depletion mutants we used for our experiments had little effect on cellular m6A levels (Figure S4K). Thus, Pho92 associates with target genes in a Paf1C-dependent manner. Our data suggest that Pho92 is loaded co-transcriptionally to nascently produced mRNAs.

### Pho92 promotes decay of m6A modified transcripts

Proteins of the YTHDF family, to which Pho92 belongs, promote mRNA decay but have also been shown to impact translation efficiency (Li *et al*, 2017; Wang *et al*., 2014; Wang *et al*., 2015; Zaccara & Jaffrey, 2020). To determine whether Pho92 plays a possible role in mRNA decay, we reasoned that if Pho92 is indeed important for the decay of m6A modified transcripts, *pho92*Δ cells should also display higher m6A levels. Hence, we determined the m6A levels by LC-MS of cells entering meiosis (4h SPO) and compared the wild type to *pho92*Δ. We found that mRNAs isolated from *pho92*Δ cells displayed a marked increase in m6A over adenosine (A) levels (Figure 5A). As expected, *ime4*Δ cells showed no detectable m6A levels.

**Figure 5.**
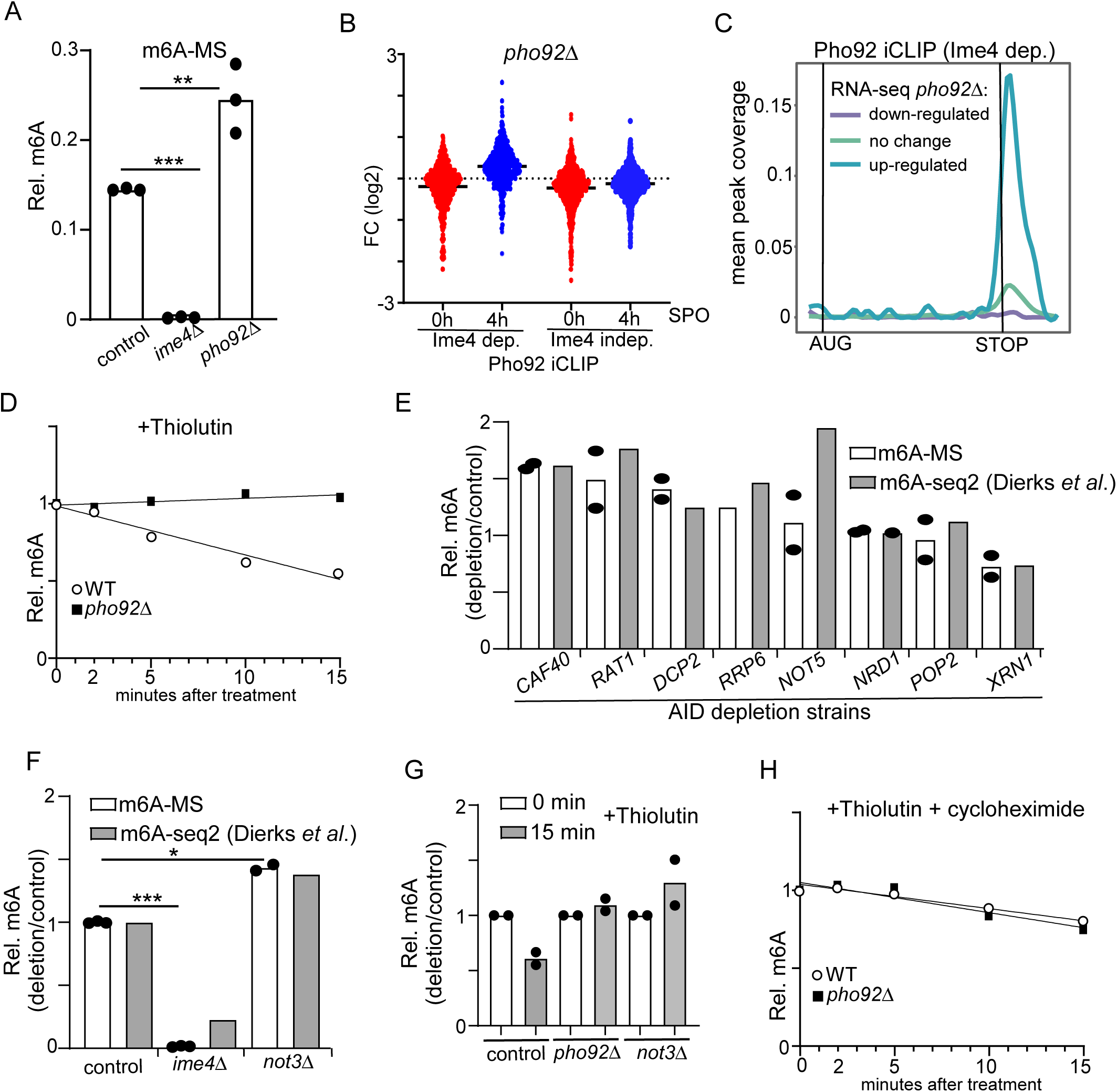
Pho92 and CCR4-NOT promotes decay of m6A marked transcripts. (**A**) m6A-MS for control, *ime4*Δ, and *pho92*Δ (FW4911, FW6060, and FW6997). Samples were taken at 6h in SPO. mRNA was isolated using oligodT paramagnetic beads. Samples were digested, and m6A levels of A were determined by LC-MS. **p< 0.01 and ***p< 0.001, compared to WT control on a two-way ANOVA followed by a Fisher’s least significant difference (LSD) test. **(B**) Differential analysis of *pho92*Δ vs WT for transcripts with Ime4-dependent and Ime4-independent Pho92 binding sites. Samples were taken at 0 and 4 hours in SPO for WT and *pho92*Δ cells (FW1511 and FW3528). RNA-seq was performed. Displayed are transcripts that are bound by Pho92 in an Ime4 dependent and independent manner as determined by iCLIP. Differential expression between 0h in SPO for *pho92*Δ vs WT are displayed (red), and 4h in SPO *pho92*Δ vs WT are displayed (blue). **(C)** Metagene analysis plotting Ime4-dependent Pho92 binding sites on genes that were either significantly upregulated, down regulated, or unchanged in 4h in SPO *pho92*Δ vs WT RNA-Seq. The y-axis represents the proportion of transcripts in that RNA-seq category with a binding site overlapping a given x coordinate. (**D**) m6A-ELISA in WT and *pho92*Δ cells (FW1511 and FW3528) after blocking transcription using thiolutin. Cells were grown and induced to enter meiosis. Cells were treated with thiolutin, and samples were taken at the indicated time points. Relative m6A levels were determined by m6A-ELISA. **(E)** Relative m6A levels in depletion mutants of various decay pathways determined by m6A-MS and m6A-seq2 (FW5958, FW6080, FW6048, FW5880, FW 6070, FW6043, FW5956, FW5964). In short, cells were grown and induced to enter meiosis. Each decay mutant bearing auxin-induced depletion alleles (AID) was depleted by adding auxin at 4h in SPO, and samples were collected at 6 hours in SPO. m6A-MS and m6A-seq2 data were normalized to a control strain harbouring the *TIR* ligase. m6A-seq2 data were obtained from Dierks et al (2021). **(F)** Similar analysis as E, except that the signals for *ime4*Δ and *not3*Δ cells are shown (control, FW4911, FW6060 and FW6093). *p< 0.05 and ***p< 0.001, for m6A-MS compared to WT control on a two-way ANOVA followed by a Fisher’s least significant difference (LSD) test. WT and *ime4*Δ data are the same as in A. (**G**) m6A-ELISA comparing WT, *pho92*Δ and *not3*Δ signal before and after blocking transcription with thiolutin for 15 min. Cells were grown to be induced to enter meiosis (FW1511, FW3528 and FW6090). Cells were treated with thiolutin, and samples were taken at 15 min after treatment. Relative m6A levels were determined by m6A-ELISA. **(H)** Similar analysis as D, but cells were treated with both thiolutin and cycloheximide.

We also performed RNA-seq (*pho92*Δ vs WT) of cells staged prior to meiosis when Pho92 is weakly expressed and m6A is relatively low (0 hours SPO) or undergoing early meiosis when Pho92 is strongly induced and m6A is high (4 hours SPO). We examined the Pho92 bound Ime4-dependent and Ime4-independent transcripts identified by Pho92 iCLIP (Figure 5B). At 0 hour in SPO, we observed little difference between *pho92*Δ and the WT (Figure 5B and S5A). At 4 hours in SPO we observed a notable increase in expression for Pho92 bound transcripts that were Ime4-dependent, but not for the Ime4 independent transcripts (Figure 5B, S5A). Out of 295 transcripts significantly up-regulated in *pho92*Δ vs WT (4 hours SPO), 95 were Pho92 targets as determined by iCLIP (Figure S5A). Moreover, transcripts significantly upregulated in *pho92*Δ at 4 hours in SPO showed greater binding of Pho92 at 3’ends compared to genes that either did not change in expression or were downregulated (Figure 5C). We observed a similar link between Pho92 Ime4-depedent transcripts and increased expression in *ime4*Δ cells during entry into meiosis (Figure S5B, S5C and S5D). For example, 212 out of 505 Pho92 iCLIP targets were significantly upregulated in *ime4*Δ vs WT (4 hours SPO) (Figure S5B). We conclude that Pho92 limits the accumulation of m6A modified transcripts.

Measuring the rate of mRNA decay of m6A transcripts is not trivial, foremost because m6A transcripts can be masked by presence of non-m6A transcripts within a pool of RNAs (Garcia-Campos *et al*., 2019). Additionally, measuring mRNA decay in sporulating cells using metabolic labelling approaches (e.g. SLAM-seq) is problematic, due to slow uptake of exogenous labelled nucleotides under these conditions (Herzog *et al*, 2017). Therefore, we adopted an alternative approach to measure decay of m6A-transcripts. We reasoned that if m6A modified transcripts are less stable compared to non-m6A mRNAs, m6A is expected to decay faster after blocking mRNA synthesis. We treated cells with thiolutin to block transcription, and subsequently determined m6A levels by m6A-ELISA or LC-MS in wild type and *pho92*Δ cells during early meiosis (4 hours SPO) (Figure 5D and S5E). Overall m6A levels declined after blocking transcription with thiolutin, indicating that m6A modified mRNAs are less stable than unmodified mRNAs. In *pho92*Δ cells however, m6A levels did not decline after blocking transcription, further supporting a role for Pho92 in promoting the decay of m6A modified mRNAs.

### CCR4-NOT mediates m6A decay

Pho92 itself has no known enzymatic activity that would drive the decay of transcripts. To identify protein complexes involved in the decay of m6A modified transcripts via Pho92, we depleted or deleted various genes involved in mRNA decay, and measured the effect on m6A levels via LC-MS in meiosis (6h SPO) (Figure 5E and 5F). Essential components were depleted using induction of the AID system at 4 hours in SPO and m6A levels were determined at 6 hours SPO (2 hours after depletion). We also compared the LC-MS data with m6A-seq2 data, a technique that relies on multiplexed m6A-immunoprecipitation of barcoded and pooled samples, of the same depletion alleles (Dierks *et al*, 2021). We found that *not3*Δ displayed an increase in m6A levels detected by LC-MS and m6A-seq2 (Figure 5F). Moreover, depletion of Caf40 and Rat1 (*CAF40-AID* and *RAT1-AID*) showed a consistent increase in m6A levels using both LC-MS and m6A-seq2 (Figure 5E). In contrast depletion of Xrn1 (*XRN1-AID*) had the opposite effect and showed reduced m6A levels. Our data suggest that the various mRNA decay pathways can have opposing effects on m6A levels during yeast meiosis. Both Caf40 and Not3 are part of the CCR4-NOT complex the major mRNA deadenylation complex in eukaryotes (Collart, 2016). Various reports have shown that YTH reader proteins facilitate recruitment of the CCR4-NOT complex to target RNAs (Du *et al*, 2016; Kang *et al*, 2014). We determined whether the CCR4-NOT complex had a similar effect on the decay of m6A modified transcripts as Pho92. We blocked transcription in *not3*Δ diploid cells entering meiosis, and measured m6A levels. We found that like Pho92, the decay of m6A-modifed transcripts was reduced in *not3*Δ diploid cells when compared to wild-type cells (Figure 5G). Thus CCR4-NOT facilitates the decay of m6A modified transcripts. Given that CCR4-NOT and Pho92 interact with each other, our data suggest that Pho92 promotes decay of its target transcripts via the CCR4-NOT complex.

### Pho92 and CCR4-NOT mediated m6A decay requires translation

Among Pho92 interactors several proteins involved in translation and ribosome biogenesis were identified (Figure 4A). In addition to the vast literature on the intimate connection between mRNA decay rate and translation elongation, a physical link between CCR4-NOT and ribosomes has also been established, implicating CCR4-NOT complex directly in translation (Buschauer *et al*, 2020). This prompted us to examine whether the decay of m6A modified transcripts relied on active translation. In addition to blocking transcription, we also inhibited translation by treating cells with cycloheximide. Blocking transcription and translation alleviated the differences in decay rates between *pho92*Δ and wild-type cells, suggesting that the decay of m6A modified transcripts is certainly dependent on translation (Figure 5H and S5E). Similarly, blocking translation and transcription alleviated the differences between *not3*Δ and wild type (Figure S5F). We conclude that Pho92 stimulates the decay of transcripts of m6A modified transcripts in a translation dependent manner and, at least in part, via the CCR4-NOT complex.

### Pho92 interacts with ribosomes

The observation that Pho92 stimulates decay of m6A marked transcripts in a translation dependent manner, suggests that Pho92 functions at ribosomes. To investigate whether Pho92 indeed associates with ribosomes, we prepared cell lysates enriched for ribosomes from cells expressing Pho92 in cycling cells by pelleting extracts with high-speed centrifugation. We observed that majority of Pho92 sedimented together with ribosomes in the pellet (Figure 6A and Figure S6A). Additionally, we performed polysome profiling to determine whether Pho92 associates with actively translating ribosomes in diploid cells undergoing early meiosis (Figure S6B). We found that Pho92 associates with polysome fractions as well as monosome fractions (Figure 6B), and hence conclude that Pho92 associates with ribosomes including translating ribosomes.

**Figure 6.**
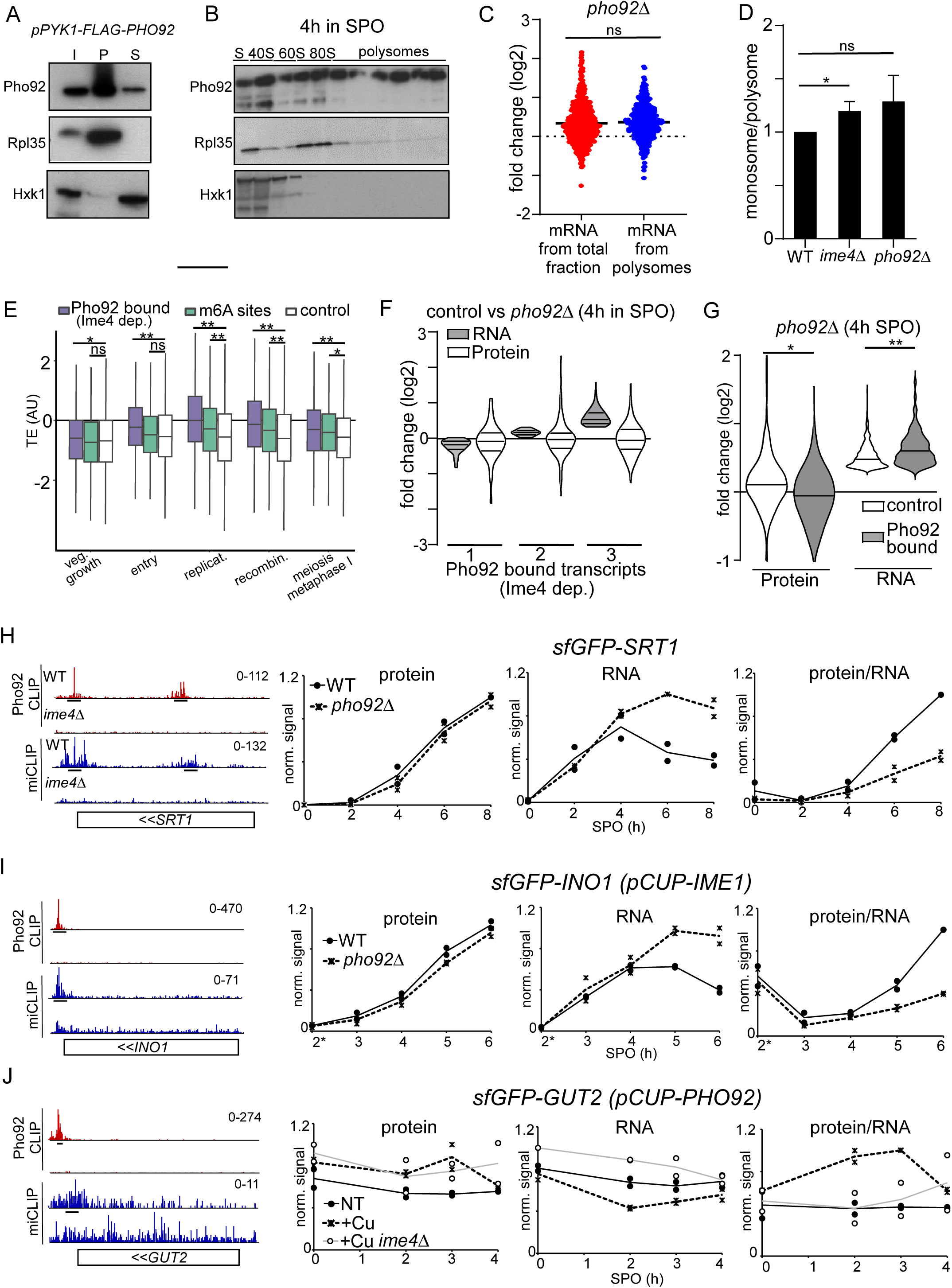
Pho92 interacts with polysomes and controls protein expression. (**A**) Sucrose cushion analysis of cells expressing *pPYK1*-FLAG-Pho92 (FW 8732) were grown in rich medium. Immunoblots of total (T), pellet/ribosomal (P) and soluble (S) fractions are shown for FLAG-Pho92, hexokinase (Hxk1) and Rpl35 (ribosomal subunit of 60S) proteins. (**B**) Polysome fractionation and western blot of Pho92. Diploid cells harbouring Pho92 tagged with V5 (FW4478). Cells were induced to enter meiosis and at 4 hours in SPO samples were taken. The small (40S), large (60S), both (80S) subunits and polysomes are highlighted from the polysome traces. Protein extracts from fractions were probed for Pho92-V5 by immunoblotting. As controls membranes were probed for Rpl35 and Hxk1. (**C**) Polyribo-seq analysis was performed 4h SPO *pho92*Δ vs WT (FW1511 and FW3528). Ribosomal fractionation was performed to isolate mRNAs bound to polysomes. Polyribo-seq was performed and the data was compared to RNA-seq from total fraction. Displayed are transcripts that are bound by Pho92 in an Ime4 dependent manner as determined by iCLIP. Differential expression between *pho92*Δ vs WT are displayed for total mRNA from total fraction (red), and polysome fraction (blue). Non significant (ns) compared to WT control on a Welsch’s t-test. (**D**) Monosome over Polysome ratio for WT (FW1511), *pho92*Δ (FW3528), and *ime4*Δ (FW725). Cells were induced to enter meiosis, samples were taken at 4 hours in SPO. From the polysome traces we determined the ratio of monosomes over polysomes. Signals for n=3 biological repeats are displayed. *p< 0.05, compared to WT control on a two-way ANOVA followed by a Fisher’s least significant difference (LSD) test. (**E**) Analysis of translation efficiency (TE) using Brar et al 2012 dataset. We assessed the TE for transcripts harbouring Ime4-dependent Pho92 binding sites, m6A sites and a control set of transcripts comprised of the rest of expressed mRNAs at 4h in SPO. Boxes show the median and interquartile range, extending lines show first and fourth quartiles. *p< 0.005 and **p< 0.001 compared to control using Welsch’s t-test. (**F**) Violin plots describing RNA-seq and proteome data from comparing *pho92*Δ (FW3528) to control (FW1511). In short, cells were induced to enter meiosis. Samples for RNA-seq and whole proteome analysis were taken at 4 hours in SPO for WT and *pho92*Δ cells. Transcripts with Ime4-dependent Pho92 binding sites and with a signal in whole proteome quantification were used for the analysis. The RNA-seq data divided in the three groups (1 to 3), while group 1 represents genes with the reduced RNA-seq signal, group 2 little change, group 3 upregulated in *pho92*Δ RNA-seq. The corresponding signal from the whole protein data is displayed. (**G**) Similar analysis as F, except that the control entailed a group of genes up-regulated in the RNA-seq that were not bound by Pho92 were used for the analysis. *p< 0.05 and **p< 0.01 compared to control on a Welsch’s t-test. **(H-J)** *SRT1, GUT2* and *INO1* transcripts bound by Pho92 and marked with m6A. Shown are the data tracks for Pho92 CLIP for WT (FW4472) and *ime4*Δ (FW4505) cells, and miCLIP for WT (FW1511) and *ime4*Δ (FW725). Underlined are the binding sites identified in the analysis. Tracks are crosslink per million normalised stranded bigWigs viewed in IGV. Relative protein, RNA, and protein over RNA ratios. WT and *pho92*Δ cells were grown to enter meiosis and samples from time points indicated. For the analysis *SRT1, INO1, and GUT2* were tagged seamlessly with sfGFP at the amino terminus. Protein expression was determined by western blotting. The relative signal is displayed with max signal for each biological repeat scaled to one. RNA levels were determined by RT-qPCR. The relative signal was computed by setting the maximum signal for each time course experiment (which included WT and *pho92*Δ) repeat to one. Right panel. The ratio of protein and RNA signals. The relative signal was computed by setting the maximum signal for each time course experiment (which included WT and *pho92*Δ) repeat to 1. For *SRT1* analysis cells were induced to enter meiosis at the indicated time points (FW9949 and FW9950). For *INO1* analysis meiosis was induced using *pCUP-IME1* synchronization system (FW9746 and FW9747). For *GUT2* analysis was performed in the presence or absence of Pho92 expression from the *CUP1* promoter (FW 10438 and *ime4Δ*, FW10441).

### Pho92 stimulates translation efficacy

Our observation that Pho92 associates with ribosomes presented the possibility that Pho92 also regulates translation. We assessed the translation efficiency of Pho92-bound transcripts by performing RNA-seq on RNA isolated from polysome fractions, known as polysome profiling (King & Gerber, 2016). We found that the Pho92 iCLIP targets were similarly upregulated in the polysome fractions compared to total RNA fraction (*pho92*Δ vs WT, 4 hours SPO), suggesting that translation efficiency as measured by polysome profiling is not directly affected in *pho92*Δ cells (Figure 6C). Interestingly, we noticed an increase in monosome over polysome ratio in *ime4*Δ cells and, though not significant, in *pho92*Δ cells, suggesting a defect in translation that was not detectable with polysome profiling (Figure 6D and Figure S6C). Moreover, we compared the translation efficiency of Pho92 targets throughout meiosis as measured by ribosome footprinting (Brar *et al*, 2012). Transcripts bound by Pho92 such as *BDF2* (Figure 1I) displayed increased ribosome footprints specifically during pre-meiotic DNA replication and recombination (Figure S6D). Meta-analysis revealed that the median translation efficiency (TE) was higher for Pho92 target transcripts (Pho92 iCLIP) and m6A modified transcripts (miCLIP) during premeiotic DNA replication, and recombination compared to a control set of transcripts (Figure 6E). These stages of meiosis also match the stages when m6A is most enriched (Schwartz *et al*., 2013). Thus, Pho92 bound transcripts have a higher translation efficiency during early meiosis and increased decay rates, which is unusual as transcripts with increased decay rates typically have a lower translation efficiency (Presnyak *et al*, 2015). Taken together, these data suggest that Pho92 in addition to regulating decay, also controls some aspect of translation.

To determine how Pho92 and m6A control protein synthesis more directly, we performed quantitative proteomics in WT and *pho92*Δ cells during early meiosis and compared the data to RNA-seq (Figure 6F and S6E). We reasoned that if Pho92 stimulates protein synthesis then in *pho92*Δ cells the protein over RNA ratio must be lower. We examined a set of genes that showed Ime4-dependent binding of Pho92 and a control set of genes. To highlight differences, we grouped the RNA-seq data (WT vs *pho92*Δ) into three categories: genes that were down-regulated (1), showed little change (2) or were up-regulated (3). For Pho92-bound transcripts, protein and RNA changes were not notably different for the group of genes that were either down-regulated (1) or showed little change in the RNA-seq (2) (*pho92*Δ vs WT) (Figure 6F and S6E). However, genes that were up-regulated (3) in the RNA-seq (*pho92*Δ vs WT), were not upregulated for the protein products (Figure 6F and S6E). In contrast, the control set of genes that were up-regulated in the RNA-seq (*pho92*Δ vs WT), showed an increase in protein levels (Figure 6F and S6E). We noticed that the increase in the RNA-seq (*pho92*Δ vs WT) of this control group was less compared to Pho92-bound transcripts. This prompted us to generate a second control set of genes that showed approximately equal increase in the RNA-seq (*pho92*Δ vs WT) but no binding of Pho92. This control group of genes also showed an increase in protein levels following the RNA-seq (*pho92*Δ vs WT), while the Pho92-bound transcripts displayed a small decrease in protein level despite the RNA levels were more increased for this set of transcripts (*pho92*Δ vs WT) (Figure 6G). We conclude that Pho92-bound transcripts with increased RNA levels were not followed by an increase in protein levels in *pho92*Δ cells, supporting the idea that Pho92 stimulates protein synthesis of its target mRNAs.

### Pho92 contributes to translation efficacy of targets

To confirm if the aforementioned trend of discordant protein and mRNA levels can also be observed for individual genes, we selected several transcripts with Ime4-dependent Pho92 binding sites to test further (Figure 6H-J and S6F-I). We tagged *INO1*, *SRT1, GUT2* at the amino-terminus seamlessly with sfGFP. Also, we tagged *RGT1* and *BDF2* at carboxy-terminus with the V5 tag. Subsequently, we performed time courses covering early meiosis from cells either directly shifted to starvation (SPO medium) or used *pCUP1-IME1* to induce meiosis in a highly synchronous manner. For each time course experiment, we determined the relative protein levels, RNA levels, and as a proxy for translation efficacy we computed the protein over RNA ratio. Consistent with the proteomics and RNA-seq data, we found that *SRT1*, *INO1*, *GUT2*, *RGT1*, and *BDF2* displayed a decrease in protein over RNA levels in *pho92*Δ cells over several time points (Figure 6H-J and S6F-I). *SRT1* showed increased RNA levels in *pho92*Δ cells compare to the control, while protein levels were higher in control cells (Figure 6H). Like *SRT1*, albeit less, the protein over RNA ratios decreased for *GUT2* and *RGT1* in *pho92*Δ cells (Figure S6G and S6H). For the analysis where *pCUP-IME1* was induced for synchronization we observed that the protein over RNA ratios of *INO1* and *BDF2* were decreased in *pho92*Δ cells compared to the control at later time points (Figure 6I, S6F and S6I).

*Vice versa*, we examined whether overexpression of Pho92 can have the opposite effect on translation efficacy. We expressed Pho92 to high levels from the *CUP1* promoter (*pCUP-PHO92*) and measured protein and RNA levels for *GUT2* (Figure 6J and S6F). In the presence of high levels of Pho92 (*pCUP-PHO92* + Cu) we observed an increase in Gut2 protein levels while RNA levels decreased, and consequently a substantial increase in protein over RNA ratio (Figure 6J and S6F). The effect was dependent on the presence of Ime4. Taken together, we conclude that Pho92 associates with ribosomes to stimulate mRNA decay and protein synthesis of m6A modified transcripts.

## Discussion

m6A modified transcripts are abundant during early yeast meiosis, yet the function of the m6A mark remains elusive. Here we show that the YTH domain containing protein Pho92/Mrb1 is likely the only m6A reader in yeast and is critical for the onset of meiosis and fitness of gametes. Importantly, we provide evidence that Pho92 co-transcriptionally associates with mRNAs to direct m6A-modifed transcripts for their translation to decay fate. Our study further shows that Pho92 promotes efficient protein synthesis of m6A modified transcripts by coupling translation efficacy to mRNA decay (Figure 7). Our work in yeast meiosis establishes a link between the previously described decay and translation functions of YTH reader proteins and m6a modified mRNAs.

**Figure 7.**
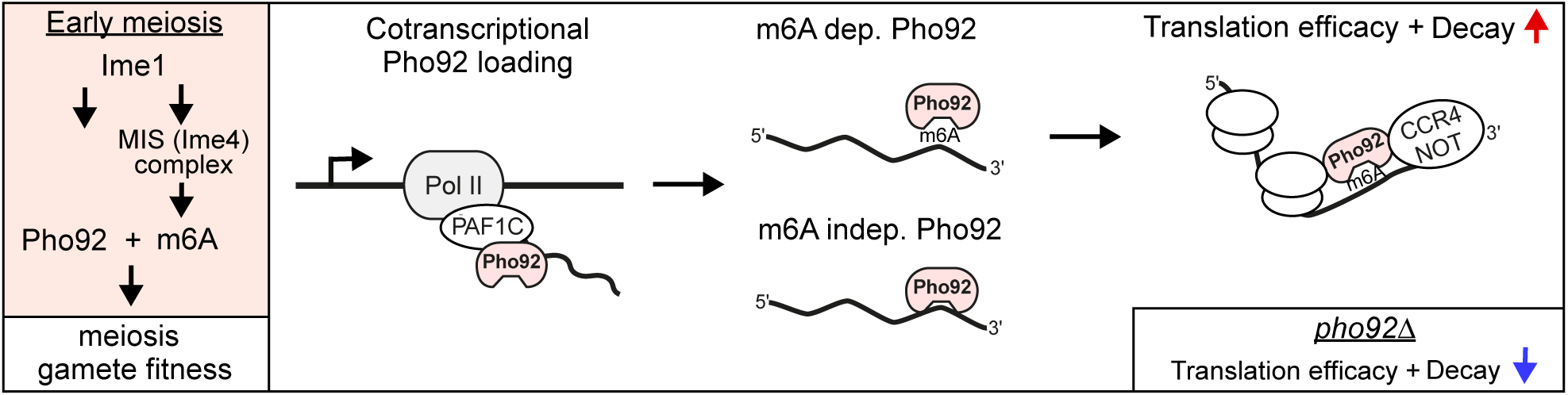
Model for role of Pho92 in early meiosis. Pho92 and MIS complex (Ime4) expression are induced by Ime1. Subsequently, Pho92 is loaded to mRNAs during transcription via Paf1C, and Pho92 promotes the translation to decay fate of m6A modified transcripts involving CCR4-NOT.

### Pho92 is key to meiotic fitness

During yeast meiosis, which is induced by severe nutrient starvation, diploid cells are triggered to undergo inarguably the most complex cell differentiation program of the yeast life cycle (Neiman, 2011; van Werven & Amon, 2011). Master transcription factor Ime1, which drives the transcription of genes in early meiosis, likely directly regulates *PHO92* transcription. Thus in yeast the m6A writer and reader machinery (Pho92 and the MIS complex) are specifically induced during early meiosis (Figure 7). Interestingly, Pho92 also regulates the stability of the *PHO4* transcript in a phosphate-dependent manner, suggesting that Pho92 may also play an additional role outside meiosis when there is no m6A (Kang *et al*., 2014).

Tight regulation of gene expression is critically important to ensure no cellular resources are wasted during meiosis. With this view, the role of Pho92 in coupling translation efficacy to mRNA decay provides an elegant strategy to facilitate increased protein synthesis and subsequent decay of mRNAs important for meiosis. One other possibility is that Pho92 is important for decay of mitotic transcripts, so that cells can enter gametogenesis, or decay of early meiotic transcripts, so that cells enter the subsequent stages of gametogenesis. If this is the case, one expects to find the m6A modified transcripts to be confined to a defined stage corresponding to either the exit from mitosis or entry into meiotic divisions. However, we observe no enrichment for genes linked to mitosis in the Pho92 iCLIP data, and m6A levels are spread over several stages during early meiosis (Schwartz *et al*., 2013). Rather, our data suggests that Pho92 increases the translation efficacy of m6A modified transcripts during critical stages of the early meiotic program when errors in meiotic chromosome segregation are fatal to the survival of gametes. Indeed, cells lacking Pho92 exhibit a delay in the onset of meiotic divisions and reduced gamete fitness.

While Pho92 deletion mutants exhibit a delay in meiosis, an Ime4 deletion mutant has more severe phenotype (Figure 3). Apart from the function that m6A has to mediate the Pho92 interaction, it is plausible that m6A has additional roles. However, it is worth noting that a catalytic dead mutant of Ime4 has a less severe phenotype in meiosis, which lead to the conclusion that Ime4 has noncatalytic function during meiosis (Agarwala *et al*., 2012). Therefore, we propose that the implications of the m6A modification is largely manifested through Pho92.

### Mechanism of co-transcriptional loading of Pho92

Our data suggests that Pho92 interacts with target transcripts during transcription via transcription elongator Paf1C (Figure 7). First, we found that Pho92 can interact with Paf1C, and that this interaction is critical for the localisation of Pho92 to the nucleus. Interestingly, the Leo1 subunit of Paf1C is also known to interact with nascent RNA, and thus forms a logical platform to facilitate the interaction between Pho92 and nascent RNA (Dermody & Buratowski, 2010). Pho92 binding sites show striking 3’ end enrichment regardless of whether binding is Ime4-independent or at m6A sites, which would suggest that the positioning is not entirely dependent on m6A. Pho92 accumulates at chromatin over the course of gene transcription, in a manner that is dependent on Paf1C. This would suggest that transcription or chromatin factors contribute towards Pho92 positioning on transcripts, which may explain why Ime4-independent Pho92 binding does not need a strong RNA sequence context. A simple mechanism could be that m6A locks Pho92 binding to RNA, which in turn allows the Pho92-mRNAs to be sent to translating ribosomes (Figure 7). It is likely that without m6A locking Pho92 in place, the protein-RNA interaction is less stable.

Transcription and chromatin mediated loading of RNA binding proteins and m6A machinery to control RNA fates is not only limited to Pho92. The co-transcriptional loading of RNA binding proteins has been reported for other RNA binding proteins to control RNA export (Fischl *et al*, 2017; Shahbabian *et al*, 2014; Viphakone *et al*, 2019). There is also evidence that the promoter sequences influence the stability of RNA (Trcek *et al*, 2011). With regard to m6A, there is proof that m6A is deposited early in RNA synthesis either co-transcriptionally or via chromatin (Huang *et al*, 2019; Ke *et al*, 2017). YTH proteins can act in the nucleus to facilitate the decay of nuclear m6A modified transcripts and thereby control chromatin states (Liu *et al*, 2020). In *S.pombe*, the YTH-RNA-binding protein Mmi1 is also reported to be co-transcriptionally recruited to ‘decay-promoting’ introns bearing sequence elements known as ‘determinants of selective removal’ (DSRs), which are enriched in meiotic mRNAs (Kilchert *et al*, 2015). Contrary to other known YTH-containing domain proteins including Pho92, Mmi1 is incapable of binding to the m6A consensus motif and the Mmi1 role is in fact reported to be suppression of activation of the meiotic programme during mitotic growth, which is in contrast to Pho92’s role in meiosis in *S. cerevisiae* (Harigaya *et al*, 2006; Shichino *et al*, 2018; Wang *et al*, 2016).

### The role of Pho92 in the translation to decay fate

Our analysis of Pho92 shows that the destiny of m6A marked transcripts in translation and mRNA decay are linked. On one hand, we have experimental evidence showing Pho92 is important for efficient protein production. Deletion of Pho92 leads to a reduction in protein over RNA ratios, suggesting translational efficacy is lowered. Additionally, Pho92 bound transcripts showed increased translation efficiency compared to a control set (Figure 6). On the other hand, Pho92 stimulates the decay of m6A modified transcripts. How does Pho92 promote decay as well as translation efficacy of m6A modified transcripts? While it is not impossible that the translation and decay functions of Pho92 are exerted via distinct mechanisms, the likelihood of these functions being linked is more plausible. Firstly, the decay of m6A marked transcripts bound by Pho92 occurred in a translation dependent manner. Secondly, we found that Pho92 associates with actively translating ribosomes, and is directly linked to increased translation efficacy. Thirdly, our analysis of decay mutants revealed that the CCR4-NOT complex facilitates the decay of m6A marked transcripts (Figure 5). Specifically, depleting the Not3 and Caf40 subunits resulted in increased m6A levels. CCR4-NOT is the major deadenylase complex in cells, which acts at ribosomes and facilitates co-translational decay (Collart, 2016). Though we did not establish a direct interaction between Pho92 and the CCR4-NOT complex in this study, others have shown that Pho92 interacts with the Pop2 subunit of CCR4-NOT (Kang *et al*., 2014).

In mammalian cells, YTHDF1 and YTHDF2 have been implicated in translation and decay, respectively (Wang *et al*., 2015; Zaccara & Jaffrey, 2020). Also YTHDC2 has been assigned both functions in decay and translation (Hsu *et al*, 2017; Kretschmer *et al*, 2018; Mao *et al*, 2019). Thus, opposite effects have been reported for YTH reader proteins. YTHDF2, the likely orthologue of Pho92, has been shown to be important for oocyte development, where it is important for decay of maternal RNAs (Ivanova *et al*, 2017). YTHDF2 associates with CCR4-NOT in mammalian cells (Du *et al*., 2016). Thus, the link between CCR4-NOT, YTH domain-containing proteins and m6A is likely conserved in yeast and mammalian gametogenesis. Interestingly, the translation function of YTHDF1 has been disputed, and it has been proposed that YTHDF proteins act redundantly to regulate decay (Zaccara & Jaffrey, 2020). Contradicting this, a significant enrichment of m6A in yeast ribosomal fractions and increased methylation of the most efficiently translated transcripts in the early stages of meiosis has been reported (Bodi *et al*., 2015). Perhaps, the decay functions of Pho92 and YTHDF1 at ribosomes facilitates translational efficiency and as consequence decay or *vice versa*. The reason that mammalian studies have failed to conclude that decay and translation functions may go hand in hand is perhaps because mammalian studies of YTH proteins have typically focused on cell lines under steady state conditions (Zaccara & Jaffrey, 2020). In contrast, yeast meiosis is a dynamic process requiring extremely quick turn over of cell RNA and protein states, which is made possible by tightly coupled RNA decay and translation.

More and more studies show that an intimate link between translation and decay exists (Pelechano *et al*, 2015; Presnyak *et al*., 2015). Slowing down translation elongation has been linked to slowing the decay of transcripts (Chan *et al*, 2018). Perhaps, Pho92 directly promotes translation elongation, which in turn leads to faster decay via the CCR4-NOT complex. As such, there is evidence that CCR4-NOT complex, apart from its direct function in translation coupled decay, monitors translating ribosomes for codon optimality (Buschauer *et al*., 2020). Another intriguing possibility is that m6A and Pho92, together with CCR4-NOT are part of an mRNA quality control mechanism to limit aberrant protein synthesis and protein misfolding, while at the same time increase productive translation. In line with this idea is that YTHDC2 protein facilitates interactions between m6A, ribosomes, and decay factors to control stability and translation of the mRNAs, and together possibly constitutes an mRNA to protein quality control mechanism (Inada, 2020; Kretschmer *et al*., 2018). How m6A, Pho92 and CCR4-NOT together promote translation and decay and how this is linked to a quality control for mRNAs and aberrant translation remains subject for further investigation.

YTH reader proteins are conserved from yeast to humans and play critical roles in developmental programs and disease pathogenesis (Patil *et al*., 2018). YTHDF1 has been shown to be involved in tumour cells immune evasion, while YTHDF2 promotes tumour cell proliferation (Chen *et al*, 2017; Han *et al*, 2019). The involvement of YTHDC1 in splicing has been associated with the incorrect processing of BRCA2, and YTHDC2 was found to contribute to metastasis (Hirschfeld *et al*, 2014; Tanabe *et al*, 2016). Additionally, YTHDF2 is a potential therapeutic target in acute myeloid leukaemia treatment (Mapperley *et al*, 2021). Understanding the molecular mechanisms of Pho92 in translation and decay in yeast may therefore reveal novel insights into human health and disease pathogenesis.

## Acknowledgements

We are grateful to the members of the laboratory of Folkert van Werven for fruitful discussions and critical reading of the manuscript. We thank members of Ule and Luscombe labs, specifically Julian Zagalak, Christoph Sadee, Patrick Toolan-Kerr, Igor Ruiz De Los Mozos, Paulo Gameiro, Federica Capraro, Andrew Steele and Flora Lee for sharing ideas, protocols and reagents towards the iCLIP and miCLIP experiments and analysis. We acknowledge Leon Chan, Elçin Ünal, and Martin Pool for sharing reagents and Schraga Schwartz and David Dierks for providing the m6A-seq2 data. We thank the Crick Advanced Sequencing, Proteomics, Metabolomics, Fermentation and Genomics Equipment Park Facilities for experimental support, specifically Phil East for helping with submission to GEO, James MacRae, Christoph Messner and Svend Kjaer for technical support. The Vermeulen lab is part of the Oncode institute, which is partly funded by the Dutch Cancer Society (KWF). This research was funded in whole, or in part, by the Wellcome Trust (FC001203, FC010110, FC001134). For the purpose of Open Access, the author has applied a CC BY public copyright licence to any Author Accepted Manuscript version arising from this submission. This work was supported by the Francis Crick Institute (FC001203, FC010110, FC001134), which receives its core funding from Cancer Research UK (FC001203, FC010110, FC001134), the UK Medical Research Council (FC001203, FC010110, FC001134), and the Wellcome Trust (FC001203, FC010110, FC001134).

## Contributions

R.A.V., and F.J.v.W. conceived the project. R.A.V., T.S. and F.J.v.W., designed the experiments. R.A.V. and T.S. generated strains, developed the miCLIP and iCLIP protocols, and performed all the biochemical, microscopy and genetic experiments with help from Z.M., A.R. and V.W.C.C. C.C. developed the data analysis pipeline and software for the miCLIP and iCLIP experiments, analysed the miCLIP and iCLIP data and integrated it with additional functional genomics datasets. I.E. and T.S. developed the m6A-ELISA protocol. E.C. and T.S. developed the m6A-MS protocol, and performed and analysed the m6A-MS experiments. I.E. analysed iCLIP data for selecting transcripts for further studies in Figure 6. H.P. analysed RNA-seq data. P.F. performed and analysed the IP-MS samples under the supervision of A.P.S. J.K. processed and analysed samples for the whole proteome-MS experiment described in Figure 6. R.R.E., performed and analysed m6A-pulldowns and dimethyl labelling MS under the supervision of M.V. R.A.V. and F.J.v.W. wrote the manuscript with input from the other authors. J.U. and N.M.L. supervised the iCLIP and miCLIP data analysis and provided funding. M.R. supervised m6A-MS and provided funding. F.J.v.W. supervised the project and provided funding.

## Additional files included

**Table S1. Yeast strains**

**Table S2. Plasmids**

**Table S3. Oligos sequences**

**Table S4. MS pulldown data**

**Table S5. iCLIP and miCLIP data table**

**Table S6. IP-MS data table Supplemental information**

**Processed RNA-seq data**

---

**Figure S1.**
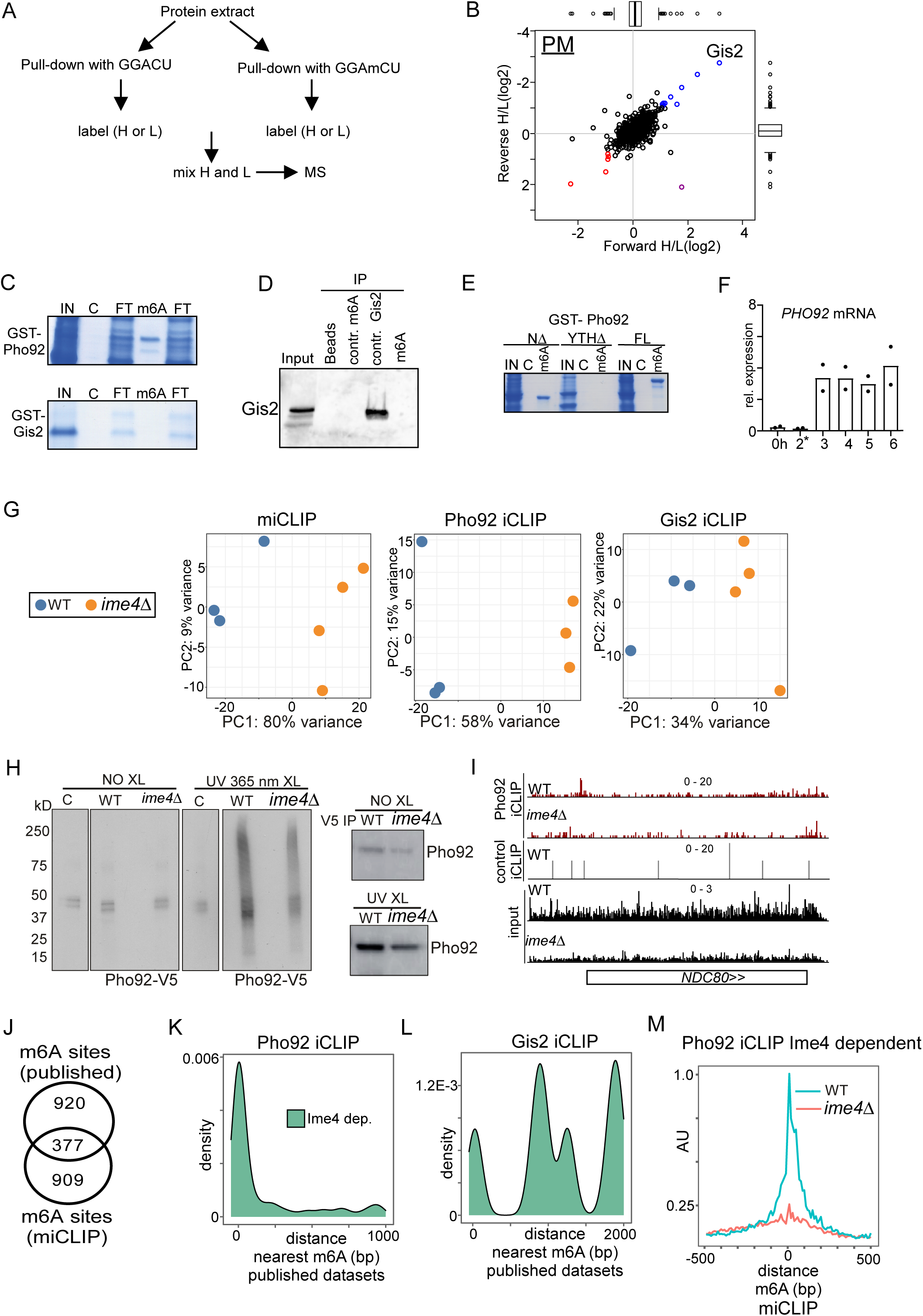
Pho92, but not Gis2, binds to m6A marked transcripts *in vitro* and *in vivo*. (**A**) Scheme describing setup for pull-down experiment. In short, protein extracts were incubated using m6A and control RNA oligo bound to streptavidin beads. Eluted proteins were labelled with heavy (H) and light (L) dimethyl isotopes, mixed, and proteins were identified by MS. (**B**) Scatter plots displaying proteins identified in m6A oligo pull down versus control in pre-meiosis (PM). (**C**) *In vitro* m6A pull down to assess binding of GST-Pho92 and GST-Gis2. Expression of GST-Pho92 and GST-Gis2 were induced in bacteria, clarified lysates were incubated with m6A consensus and control RNA oligos bound to streptavidin beads, and eluates were run on an SDS-page gel and Coomassie-stained. Input (IN), Unlabelled oligo (C), m6A oligo (m6A), Unbound flow through after incubation (FT). (**D**) Gis2 does not bind to m6A labelled or unlabelled oligos but does to control RNA oligos. Protein extracts expressing Gis2-V5 (FW 3312) were incubated with unlabelled oligo, m6A oligo, and control oligo harbouring the canonical Gis2 motif (GA(A/U)). (**E**) Similar as C except that lysates from truncations of GST-Pho92 in N-terminus (NΔ) and YTH domain (YTHΔ) were used. (**F**) Pho92 expression prior to and after induction of Ime1 expression. Cells harbouring *pCUP-IME1* and Pho92 tagged with V5 (FW 9962) were induced to enter meiosis in sporulation medium (SPO). After 2 hours in SPO, Ime1 was induced with copper sulphate (labelled with *). Samples were taken at the indicated time points, and RNA levels were determined by RT-qPCR. *ACT1* was used for normalization. (**G**) Principle component analysis (PCA) of miCLIP (left), Pho92 iCLIP (middle), and Gis2 iCLIP (right) at the level of counts per peak. Indicated are the biological repeats for the WT (blue) and *ime4*Δ (yellow). (**H**) Autoradiograph showing the protein RNA complexes in control untagged (C) and Pho92-V5 cells. In short, cells were grown till 4 hours in SPO in presence of 4-thiouracil. Cell were harvested and either crosslinked or left untreated. Protein extracts were generated, and Pho92 was immunoprecipitated with anti V5 antibodies. RNA-protein complexes were labelled with (γ-^32^P)-ATP, and separated by SDS page, and transferred to nitrocellulose membrane. Displayed are the signals obtained for untagged control (FW 4256), Pho92-V5 (WT, FW 4472), and Pho92-V5 in *ime4*Δ (FW 4505). In right panel, western blots probed with anti-V5 showing the immunoprecipitation of Pho92. (**I**) Integrative genome browser (IGV) tracks of *NDC80* gene for Pho92 CLIP in WT and *ime4*Δ, and untagged control CLIP in WT background. Tracks are crosslink per million normalised, strand-specific bigWigs. (**J**) A Venn diagram showing the number of miCLIP and published m6A sites within 100nt of each other. (**K**) Metanalysis comparing the distance between Ime4-dependent Pho92 binding sites to curated m6A sites from published datasets. For each Pho92 binding site the distance to the nearest m6A site was calculated. (**L**) Similar as K except that Gis2 Ime4 dependent sites were compared to the curated m6A sites. (**M**) Metagene analysis of Pho92 CLIP Ime4 dependent sites for WT and *ime4*Δ cells. Data was centred on the m6A sites identified with miCLIP.

**Figure S2.**
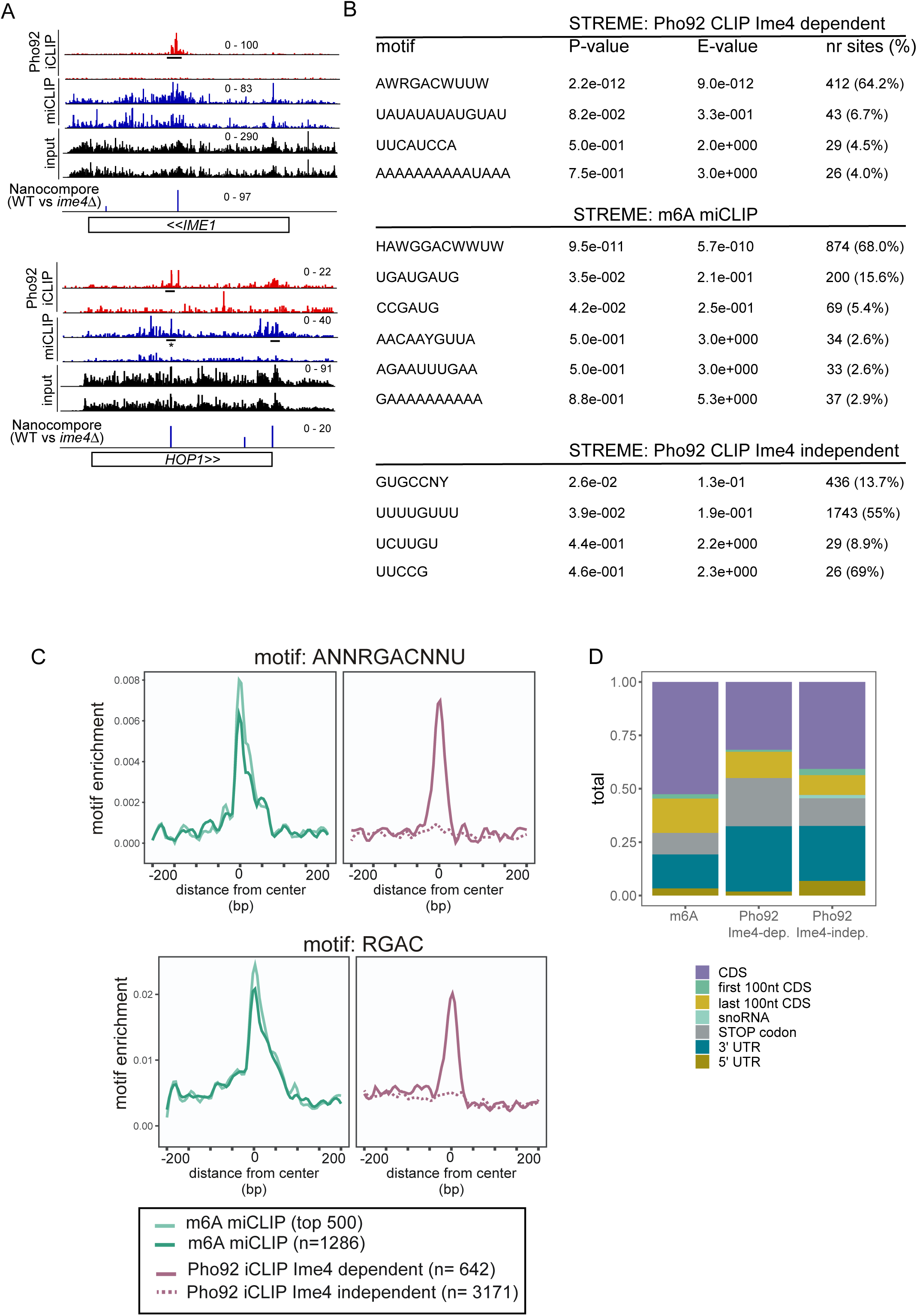
Features of transcripts bound by Pho92. (**A**) *IME1* and *HOP1* IGV tracks. Shown are the crosslinks per million normalised bigWigs for Pho92 iCLIP, and miCLIP in WT and *ime4*Δ cells. Underlined are the binding sites identified in the analysis. * this site was reported in our analysis below the significance threshold (miCLIP). The bottom IGV track shows the significant m6A sites identified with comparative Nanopore direct RNA sequencing, with the scale of the bars representing -log10(p value) from a logistic regression test, (Nanocompore, WT versus *ime4*Δ) (Leger *et al*, 2021). **(B)** Motif analysis of Ime4-dependent Pho92 binding sites, m6A sites (miCLIP) and Ime4-independent Pho92 binding sites using STREME. Significant motif sequences are shown. **(C)** Motif enrichment around m6A sites and Pho92 Ime4-dependent and independent binding sites for RGAC and ANNRGACNNU motifs. Motif frequency plotted around the centre of Pho92 binding sites, split into Ime4-dependent sites (solid line) and Ime4-independent sites (dotted line). (**D**) Stacked bar graph showing peak distributions over transcript regions for m6A sites, and Pho92 Ime4-dependent and independent binding sites.

**Figure S3.**
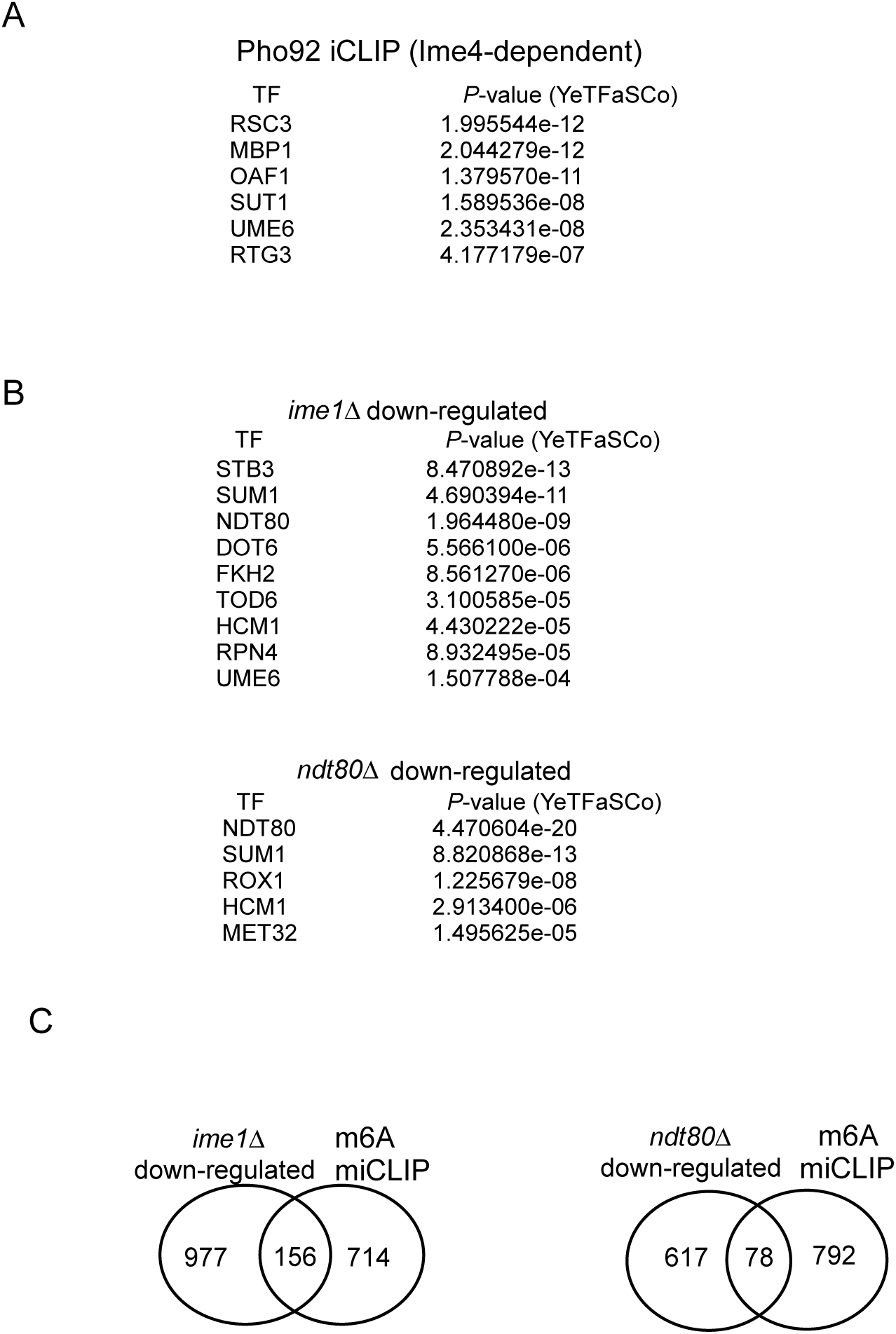
Pho92 is important for meiosis and fitness of gametes. (**A**) Motif analysis of promoters of transcripts bound by Pho92 in an Ime4-dependent manner. YTFaSCo was used for the analysis (de Boer & Hughes, 2012). (**B**) Motif analysis of promoters down-regulated in *ime1*Δ (FW81) and *ndt80*Δ (FW 4911) cell during early meiosis. YTFaSCo was used for the analysis (de Boer & Hughes, 2012). (**C**) Venn diagram displaying the comparison between RNA-seq of *ime1*Δ (FW81) and *ndt80*Δ (FW4911), and transcripts determined by miCLIP. For this analysis, Ime1 and Ndt80-dependent genes were selected by taking the transcripts that were significantly down-regulated compared to the WT control in the *ime1*Δ and *ndt80*Δ.

**Figure S4.**
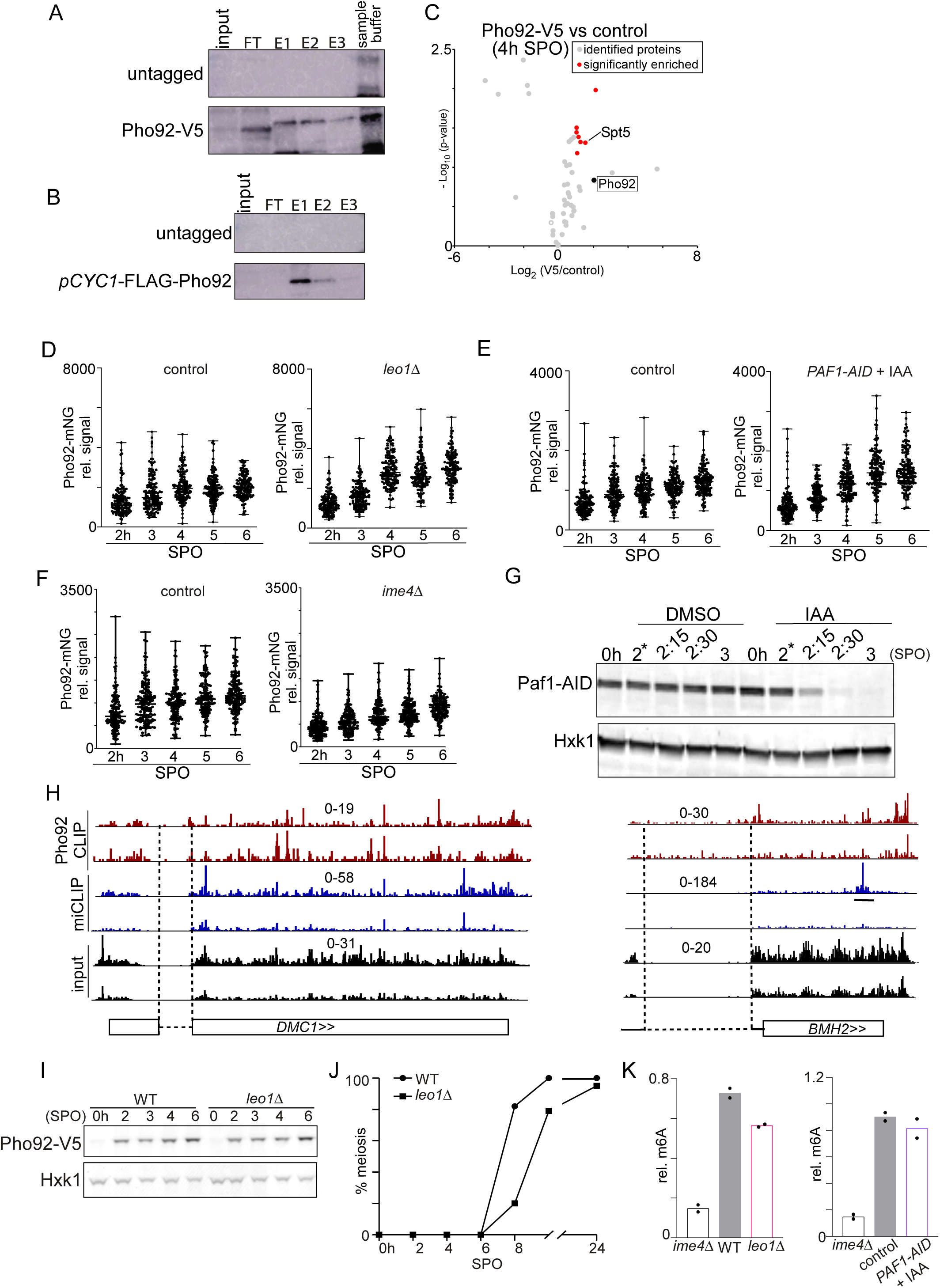
Paf1C interacts with Pho92 to direct Pho92 to nucleus. (**A**) Western blot of Pho92 eluates used for MS analysis. Diploid cells harbouring Pho92-V5 tagged and untagged control (FW4478 and FW1511). Cells were grown and induced to enter meiosis, and samples were collected at 4h in SPO. Cell extracts were incubated with anti-V5 beads and eluted with V5 peptides followed by laemmli sample buffer boiling elution. E1, E2 and E3 represent peptide eluates. (**B**) Similar as A, except that cells with the *CYC1* promoter (*pCYC1*-FLAG-Pho92, FW8734) controlling Pho92 expression and a FLAG-tag at the amino terminus was grown in rich medium conditions. Cell extracts were incubated with anti-FLAG beads and eluted with FLAG peptides. E1, E2 and E3 represent eluates. **(C)** Volcano plots of IP-MS of Pho92-V5 compared to untagged control. Significantly enriched proteins are displayed in red. Pho92 is labelled in black. (**D**) Whole cell quantification of Pho92-mNG in WT and *leo1*Δ cells during entry into meiosis. These cells also harboured *pCUP-IME1*. Cells were treated with copper sulphate at 2 hours. Samples were taken at the indicated time points. Signal for Pho92-mNG in WT and *leo1*Δ (FW9633 and FW9736). At least 150 cells were quantified per time point. (**E**) Similar as E, except that Paf1 depletion allele was used for the analysis. Paf1 fused to the auxin induced degron (AID) was used (FW10128). These cells expressed *TIR1* ligase under control of the *CUP1* promoter. The depletion was induced by treating cells with copper sulphate and IAA. Cells harbouring the *TIR1* ligase alone were used as controls (FW10129). (**F**) Same as in D, except that *ime4*Δ cells (FW9604) were used for the analysis. **(G)** Western blot of strains used in F showing the depletion of *PAF1-AID*. * indicates treatment with copper sulphate and IAA. (**H**) Pho92 binds to some intronic regions in transcripts. IGV data tracks of Pho92 iCLIP, miCLIP, and input in WT and *ime4*Δ cells. Intron regions of *BMH2* and *DMC1* are shown. (**I**) Pho92 expression in WT and *leo1*Δ cells during entry into meiosis. Diploid cells harbouring Pho92-V5 in WT and *leo1*Δ (FW4478 and FW10113) were induced to enter meiosis. Samples were taken at the indicated time points, and protein extracts were assessed by western blotting with anti-V5 antibodies. As a loading control Hxk1 was used. (**J**) Onset of meiosis in WT and *leo1*Δ cells (FW4478 and FW10113). Cells were shifted to SPO, and samples were taken at the indicated time points. Cells were fixed, and stained, DAPI masses were counted for at least n=200 cells per biological repeat. Cells with two or more DAPI masses were considered as undergoing meiosis. (**K**) m6A ELISA during entry into meiosis of WT, *ime4*Δ and *leo1*Δ cells (left) and after Paf1 depletion as described in E (right). The relative m6A signals are displayed.

**Figure S5.**
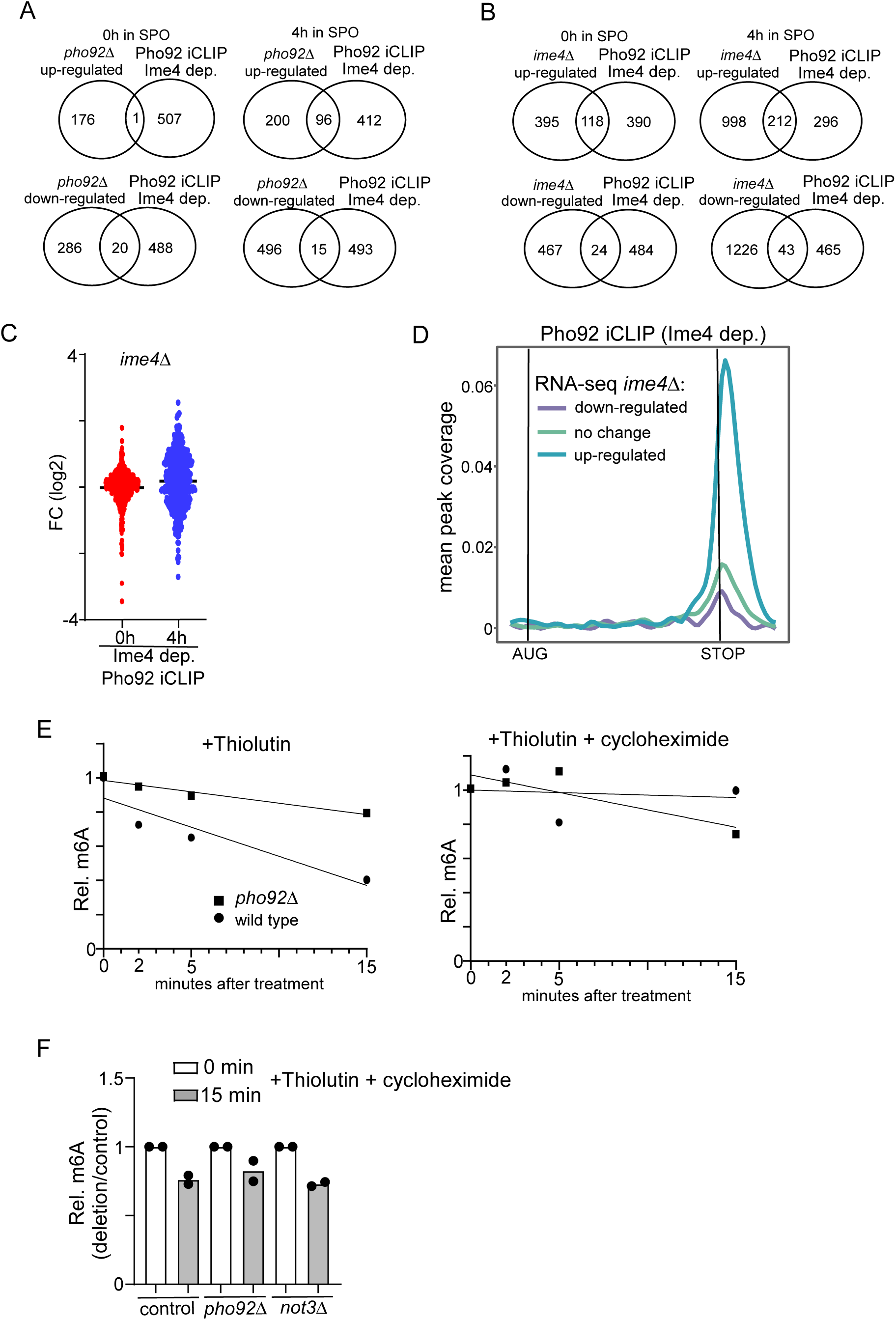
Pho92 and CCR4-NOT promotes decay of m6A marked transcripts. (**A**) Venn diagrams comparing differentially expressed transcripts in *pho92*Δ RNA-seq vs transcripts with Ime4-dependent Pho92 binding. (**B**) Similar as A, except that differentially expressed transcripts from *ime4*Δ vs WT RNA-seq is compared to Pho92-bound transcripts. (**C**) Differential analysis of *ime4*Δ vs WT for transcripts with Ime4-dependent Pho92 binding sites. Samples were taken at 0 and 4 hours in SPO for WT and *pho92*Δ cells (FW1511 and FW725). RNA-seq was performed. Displayed are transcripts that are bound by Pho92 in an Ime4 dependent manner as determined by iCLIP. Differential expression between 0h in SPO for *ime4*Δ vs WT are displayed (red), and 4h in SPO *ime4*Δ vs WT are displayed (blue). (**D**) Metagene analysis plotting Ime4-dependent Pho92 binding sites on genes that were either significantly upregulated, down regulated, or unchanged in 4 hours SPO *ime4*Δ vs WT RNA-seq. The y-axis represents the proportion of transcripts in that RNA-seq category with a binding site overlapping a given x coordinate. (**E**) m6A-MS in WT and *pho92*Δ cells (FW1511 and FW 3528) after blocking transcription using thiolutin. Cells were grown and induced to enter meiosis. Cells were treated with thiolutin (left panel) or thiolutin + cycloheximide (right panel), and samples were taken at the indicated time points. Relative m6A levels were determined by LC-MS. **(F)** m6A-ELISA comparing WT, *pho92*Δ and *not3*Δ signal before and after blocking transcription with thiolutin for 15 min. Cells were grown to be induced to enter meiosis (FW1511, FW3528 and FW6090). Cells were treated with thiolutin + cycloheximide, and samples were taken at 15 min after treatment. Relative m6A levels were determined by m6A-ELISA.

**Figure S6.**
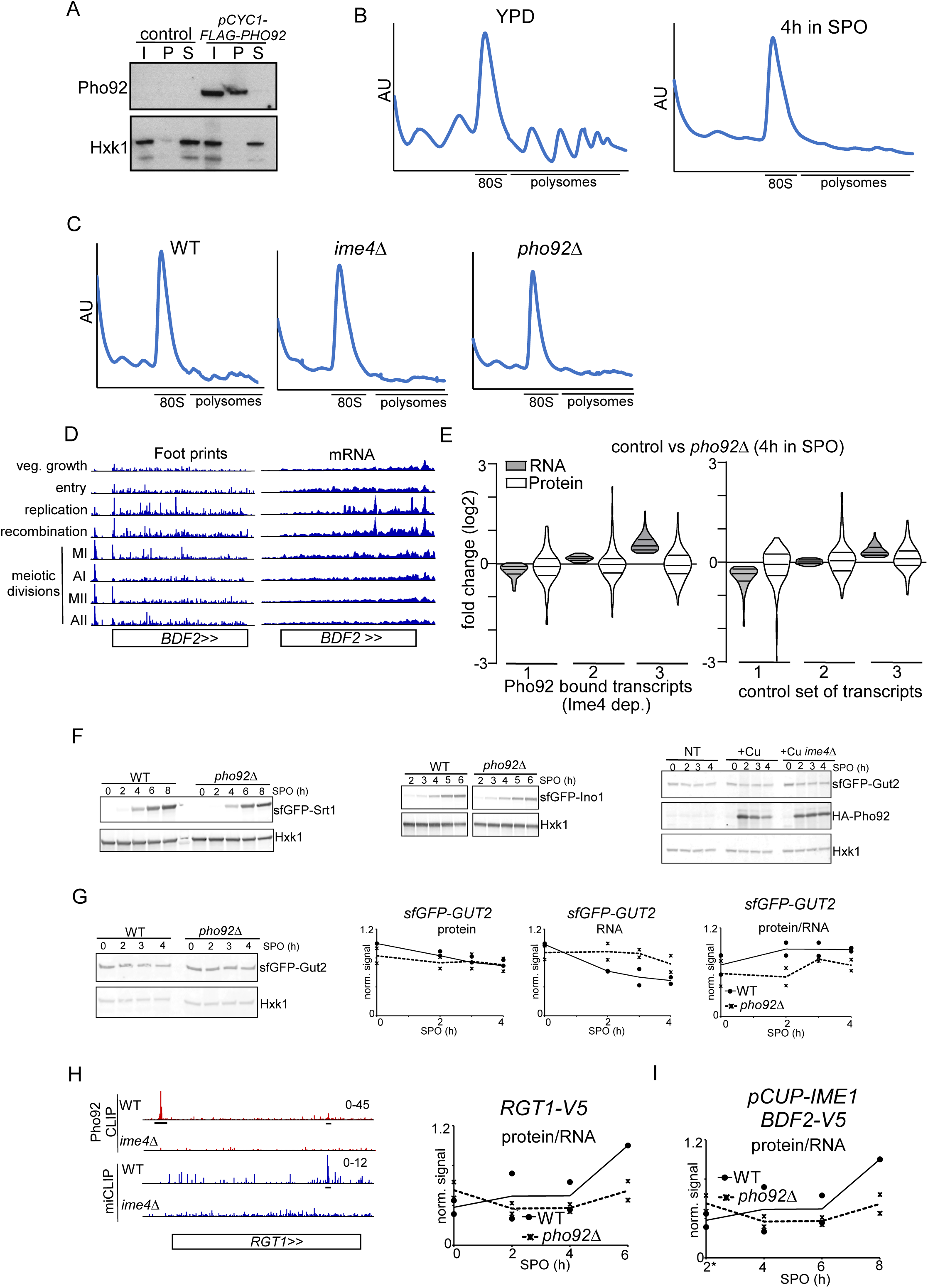
Pho92 interacts with polysomes and controls protein expression. (**A**) Sucrose cushion of cells expressing *pCYC1*-FLAG-Pho92 (FW 8734). Cells were grown in rich medium. Immunoblots of total (T), pellet/ribosomal (P) and soluble (S) fractions are shown for FLAG-Pho92, hexokinase (Hxk1) and Rpl35 (ribosomal subunit of 60S) proteins. (**B**) Polysome traces from cells grown in rich medium (YPD), and cells in early meiosis (4h in SPO). (**C**) Polysome traces of cells entering meiosis (SPO 4H) for WT (FW1511), *ime4*Δ (FW5486), and *pho92*Δ (FW3528) cells. (**D**) Ribosome footprint and mRNA-seq (RPKM) data from Brar et al (2012) at the *BDF2* locus. *BDF2* has multiple m6A peaks and Pho92 binding sites. Indicated are the different phases of yeast gametogenesis. **(E)** Related to Figure 6F. A control set of transcripts was included for the analysis that showed no binding to Pho92. **(F)** Related to Figure 6H-J, sfGFP-Srt1, sfGFP-Ino1, and sfGFP-Gut2 protein expression was determined by western blot. **(G)** Similar analysis as 6H, except that *GUT2* protein and RNA signals are displayed. For the analysis *GUT2* was seamlessly tagged with sfGFP at the N-terminus (FW9836 and FW9781). **(G)** Similar analysis as 6H, except that *GUT2* protein and RNA signals are displayed. For the analysis *GUT2* was seamlessly tagged with sfGFP at the N-terminus (FW9836 and FW9781). **(H)**). Similar analysis as G, except that *RGT1* protein over RNA signals are displayed. For the analysis *RGT1* was tagged with V5 at the C-terminus (FW9951 and FW9952). Also shown are the *RGT1* tracks for Pho92 iCLIP for WT (FW4472) and *ime4*Δ (FW4505) cells, and miCLIP for WT (FW1511) and *ime4*Δ (FW725). Underlined are the binding sites identified in the analysis. Tracks are crosslink per million normalised stranded bigWigs viewed in IGV. **(I)** Similar analysis as G, except that *BDF2* protein over RNA signals are displayed. For the analysis meiosis was induced using *pCUP-IME1* synchronization system. For the analysis *BDF2* was tagged with V5 at the C-terminus (FW 8973 and FW 9093).

