## Supplementary material for "m6A reader Pho92 is recruited co-transcriptionally and couples translation efficacy to mRNA decay to promote meiotic fitness in yeast": methods

### Plasmids and yeast strains

Yeast strains used in this study were derived either from the sporulation proficient SK1, or poor sporulation S288C or A364A backgrounds. All experiments were carried out in diploid cells apart from the experiments using strains expressing N-terminal Flag tagged *PHO92* from *PYK1* or *CYC1* promoters.

GST tagged Pho92 full length, truncations NTDΔ (containing residues 141-306) and YTHΔ (containing residues 1-140) and GST-tagged Gis2 were generated by Gibson cloning using yeast genomic DNA as template. Plasmids and oligonucleotide sequences are listed in the Tables S2 and S3 respectively.

Gene deletions and epitope tagging were achieved using the one-step deletion protocol as described previously (Longtine *et al*, 1998). Pho92 and Gis2 were tagged with copies of the V5 epitope tag using tagging cassette described in (Chia *et al*, 2017). *PAF1* was depleted using C-terminal auxin-inducible degron (AID) tag as described by (Nishimura *et al*, 2009). For efficient depletion during meiosis we used copper inducible *Oryza sativa* TIR (*osTIR*) ubiquitin ligase under the control of the *CUP1* promoter (Chia *et al*, 2021). The AID depletions of decay factors (*CAF40*, *RAT1*, *DCP2*, *RRP6*, *NOT5*, *NRD1*, *POP2*, *XRN1*) were described previously (Dierks *et al*, 2021). Seamless N-terminal tagging with superfolder GFP was performed as previously described (Khmelinskii *et al*, 2011). *URA3* gene served as a selection marker for positive selection in medium lacking uracil and counter selection in medium containing 5-fluoroorotic acid (5-FOA). Pho92-mNeonGreen fusions were constructed by tagging Pho92 with mNeonGreen tagging cassettes (Arguello-Miranda *et al*, 2018). Cells expressing Pho92-mNeonGreen fusions also harboured histone H2B fused with mCherry to determine nuclear localization. N- terminal 3X Flag- tagged Pho92 constitutive expression in S288C was achieved using the plasmids described in (Zhang *et al*, 2017). The strain genotypes are listed in Table S1.

**Growth conditions**

All experiments were performed at 30°C in a shaker incubator at 300 rpm. Cells were grown in YPD [1.0% (w/v) yeast extract, 2.0% (w/v) peptone, 2.0% (w/v) glucose, and supplemented with uracil (2.5 mg/l) and adenine (1.25 mg/l)].

For obtaining exponentially growing S288C cells for co-immunoprecipitations, yeast cells were grown in YPD to saturation overnight, diluted to $OD_{600}$ = 0.2 and subsequently were harvested after two to three doublings.

To induce entry into meiosis in SK1, a standard protocol for sporulation was followed. Cells were grown till saturation for 24h in YPD, then diluted at $OD_{600}$ = 0.4 to pre-sporulation medium BYTA [(1.0% (w/v) yeast extract, 2.0% (w/v) bacto tryptone, 1.0% (w/v) potassium acetate, 50 mM potassium phthalate] grown for about 16 h, subsequently centrifuged, washed with sterile milliQ water, centrifuged again, re-suspended at $OD_{600}$ = 1.8 (with an exception for all CLIP experiments where cells were resuspended at $OD_{600}$ = 2.5) in sporulation medium (SPO) [0.3% (w/v) potassium acetate and 0.02% (w/v) raffinose)] and incubated at 30°C. Cells were collected for protein, RNA, and DAPI staining analyses at the desired meiotic time points via pelleting at 1500g, 2min. To induce meiosis in S288C and A364A, cells were moved from saturated YPD to SPO medium and samples collected for DAPI counting every 24 hours until 72 hours.

To initiate sporulation synchronously from SK1 cells expressing *IME1* from *CUP1* promoter, 50 µM $CuSO_4$ was added to the medium 2 hours after shifting them to SPO. In cells expressing *NDT80* from *GAL1* promoter (*pGAL-NDT80*) and the transcription factor *GAL4*-ER, 1 µM β-estradiol was added at the same time $CuSO_4$.

For the m6A pull down experiment, synchronous sporulation was induced following the method previously described (Chia and van Werven, 2016). Approximately 0.05 OD of exponentially growing yeast were inoculated into reduced glucose YPD . Cells were cultured overnight and washed in milli Q water before suspending them in SPO medium at $OD_{600}$ of 2.5. After 2 hours, 50 µM of $CuSO_4$ was added to induce expression from the *CUP1* promoter and initiate sporulation synchronously.

To enable efficient Paf1 depletion (*PAF1-AID*) for microscopy experiments, 1 mM of indole-3-acetic acid (IAA) and 50 µM of $CuSO_4$were added 2 hours after cells were shifted to SPO. For mRNA decay machinery depletion mutants, IAA and $CuSO_4$were added at 4h after shifting the cells to SPO and samples collected at 6h in SPO. As control, same volume of dimethyl sulphoxide (DMSO) was added to yeast cells.

**DAPI counting and spore viability**

Cells were collected from sporulation cultures, pelleted via centrifugation (1500g, 2min), and fixed in 80% (v/v) ethanol for a minimum of 2h at 4°C. The cells were pelleted again (1500g, 2min) and resuspended in PBS with 1 µg/ml DAPI. The proportion of cells containing one or multiple nuclei were counted using a fluorescent inverted microscope.

Tetrad/ dyad counts S288C and A364A: WT and *pho92Δ* strains were patched on YPD plates and incubated for 2 days. The patches were transferred on SPO plate and further incubated for 1 week at room temperature. Colonies were suspended in 10µL of water and observed under 40x objective (Axiostar Plus (Zeiss). Number of packaged spores per ascus were counted for 200 asci.

Spore viability SK1: Diploid cells were patched from YPD agar plates to SPO agar plates and incubated for 3 days at 30°C or 37°C. Subsequently at least 40 tetrads (160 spores) were dissected and incubated at 30°C for 72h on YPD agar plates before spore survival was assessed.

**RNA extraction**

Yeast cells were harvested for RNA extraction by centrifugation, then washed once with sterile water prior to snap-freezing in liquid nitrogen. RNA was extracted from frozen yeast cell pellets using Acid Phenol:Chloroform pH 4.5 and Tris-EDTA-SDS (TES) buffer (0.01 M Tris-HCl pH 7.5, 0.01 M EDTA, 0.5% w/v SDS), by rapid agitation (1400 RPM, 65 °C for 45 minutes). After centrifugation, the aqueous phase was obtained and RNA was precipitated at -20 °C overnight in ethanol with 0.3 M sodium acetate. After centrifugation and washing with 80% (v/v) ethanol solution, dried RNA pellets were resuspended in DEPC-treated sterile water. Samples were further treated with rDNase (cat no 740.963, Macherey-Nagel) and column purified (cat no 740.948, Macherey-Nagel).

**RT-qPCR**

For reverse transcription, ProtoScript II First Strand cDNA Synthesis Kit (New England BioLabs) was used and 500 ng of total RNA was provided as template in each reaction. qPCR reactions were prepared using Fast SYBR Green Master Mix (Thermo Fisher Scientific) and transcript levels were quantified from the cDNA on Quantstudio 7 Flex Real Time PCR instrument. Signals were normalised over *ACT1*. Oligo sequences are listed in Table S3.

**RNA-Seq**

For RNA sequencing, 1 µg of total yeast total RNA was used. Libraries were prepared using the KAPA RNA hyperPrep kit (KK8540, Roche) according to the manufacturer's instructions. Libraries were sequenced on an Illumina HiSeq 4000 to an equivalent of 75 bases single-end reads, at a depth of approximately 20 million reads per library.

Adapter trimming was performed with cutadapt (version 1.9.1) (Martin, 2011) with parameters "--minimum-length=25 --quality-cutoff=20 -a AGATCGGAAGAGC -A AGATCGGAAGAGC". BWA (version 0.5.9-r16) (Liao *et al*, 2014) using default parameters was used to perform the read mapping independently to both the *S. cerevisiae* (assembly R64-1-1, release 90) genome. Genomic alignments were filtered to only include those that were primary, properly paired, uniquely mapped, not soft-clipped, maximum insert size of 2kb and fewer than 3 mismatches using BamTools (version 2.4.0; (Barnett *et al*, 2011)). Read counts relative to protein-coding genes were obtained using the featureCounts tool from the Subread package (version 1.5.1) (Liao *et al.*, 2014). The parameters used were "-O --minOverlap 1 --nonSplitOnly --primary -s 2 -p -B -P -d 0 -D 1000 -C --donotsort".

Differential expression analysis was performed with the DESeq2 package (version 1.12.3) within the R programming environment (version 3.3.1) (Love *et al*, 2014). An adjusted p-value of <= 0.01 was used as the significance threshold for the identification of differentially expressed genes.

**Western blotting**

3.6 ODs of yeast cells were pelleted from cultures via centrifugation (1500g, 2min) for protein expression analysis. Proteins were precipitated from whole cells via trichloroacetic acid (TCA) incubation for a minimum of 2h at 4°C. TCA was completely removed from the pellet via an acetone wash. Cells were lysed with glass beads using a bead beater and lysis buffer (50 mM Tris, 1 mM EDTA, and 2.75 mM DTT). Proteins were denatured in 3X sample buffer (9% (w/v) SDS and 6% (v/v) β-mercaptoethanol) at 100°C and separated by SDS/PAGE (4-12% gels). A PVDF membrane was used for protein transfer. Blocking was performed using 5% (w/v) dry skimmed milk. Proteins were detected using an ECL detection kit or using infra-red fluorescent antibodies visualised with LiCor CLx.

**Recombinant protein expression and purification**

BL-21 DE3 were used for GST-fusion protein expression. Cells were grown in TB medium supplemented with 100 µg/mL ampicillin. IPTG was added to a final concentration of 0.5 mM, after which protein expression was induced for 4 h at 30 °C. Cells were then harvested, and resuspended in lysis buffer (50 mM Tris-HCl, pH 8.0, 15% glycerol, 1 mM EDTA, 0.5 mM PMSF, 1 mM DTT, 100 mM NaCl, 1% Triton

X-100, 0.25% NP-40, protease inhibitor cocktail, and lysozyme to a final concentration of 0.5 mg/ml). Cells were lysed by freeze-thaw cycles and short sonication. Bacterial debris was removed by ultra-centrifugation at 55,000 r.p.m. for 60 min at 4 °C, after which soluble extracts were aliquoted and snap-frozen in liquid nitrogen until further use.

**Dimethyl-labelling based m6A RNA pulldowns**

RNA pull downs were performed using unlabelled lysates followed by dimethyl labelling essentially as described in (Edupuganti *et al*, 2017). RNA probes used for this experiment are prepared in the lab of Thomas Carell (Edupuganti *et al.*, 2017) and listed in Table S3. 40 µl of suspended beads (Dynabeads M280 Streptavidin) for two reactions (heavy and light, 20 µl per pull down) were first washed twice in 1 ml RNA binding buffer (50 mM HEPES-HCl, pH 7.5, 150 mM NaCl, 0.5% NP40 (v/v), 10 mM $MgCl_2$) and then incubated with RNase inhibitor RNasin plus (Promega) in RNA binding buffer (100 µl buffer with 0.8 units of RNasin/µl) for 15 minutes on ice. After removal of the buffer, beads were preblocked with yeast tRNA (100 µg/mL; AM7119; Life Technologies) in RNA binding buffer for one hour at 4 °C on a rotation wheel. The preblocked beads were washed twice with RNA binding buffer and then incubated with 5 µg of biotinylated RNA probe (per pulldown) diluted with RNA binding buffer to a final volume of 600 µl for 30 min at 4 °C in a rotation. The beads were washed once with 1 mL of RNA wash buffer (50 mM HEPES-HCl, pH 7.5, 250 mM NaCl, 0.5% NP-40 and 10 mM $MgCl_2$) and twice with protein incubation buffer (10 mM Tris-HCl, pH 7.5, 150 mM KCl, 1.5 mM $MgCl_2$, 0.1% (v/v) NP-40, 0.5 mM

DTT, and complete protease inhibitors (HALT)). During the last wash, beads in each vial, were split into two tubes one each for light and heavy labelling.

Beads containing immobilized RNA were then incubated with 500 ug of meiotic yeast extracts each in a total volume of 600 µl of protein binding buffer. The incubation reaction also contained 30 µg of yeast tRNA to prevent nonspecific binding, and RNasin. The reactions were incubated at room temperature for 30 min and then for 90 min on a rotation wheel at 4 °C. The beads were then washed three times with protein incubation buffer and twice with ice-cold PBS to remove detergent from the beads.

Proteins were on-bead digested with trypsin. Briefly, beads were resuspended in 100 µl of elution buffer (50 mM Tris, pH 8.5, 2 M urea and 10 mM DTT) and then incubated for 20 min at room temperature in a thermoshaker at 1,100 r.p.m. Iodoacetamide was then added to a final concentration of 55 mM, and the mixture was incubated for 10 min in a thermoshaker (1,100 r.p.m.) at room temperature in the dark. Proteins were then partially digested from the beads by the addition of 250 ng of trypsin for 2 h at room temperature in a thermoshaker in the dark. After incubation, the supernatant was collected in a separate tube. The beads were then incubated with 50 µL of elution buffer for 5 min at room temperature in a thermoshaker (1,100 r.p.m.). 100 ng of fresh trypsin was added to the pooled eluates, and proteins were digested overnight at room temperature.

Tryptic peptides from individual pulldowns obtained after on-bead digestion were differentially labelled with dimethyl isotopes ($CH_2O$ or $CD_2O$) essentially as described (Boersema *et al*, 2009). Forward and reverse reactions were set up by mixing labelled peptides as follows: Forward reaction: control probe, $CH_2O$ (light); m6A probe, $CD_2O$ (medium) and Reverse reaction: control probe, $CD_2O$ (medium); m6A probe, $CH_2O$ (light). The reverse experiment represented a biological-replicate label swap. Labelled reactions were acidified to pH <2 with TFA (10%) and desalted with C18 Stage tips before MS analyses on the Orbitrap Fusion Tribrid mass spectrometer (Thermo), a similar LC gradient was used, and samples were measured in top-speed mode. Data processing was done with the MaxQuant software. Raw data were analyzed with DimethylLys0 and DimethylLys4 (light and medium) labels and matching between runs was enabled. UniProt database for *S. cerevisiae* downloaded on 13 July 2014 was used for identification. To identify significant interactors, normalized ratios from MaxQuant output tables for the forward and reverse pulldowns were plotted.

For validation of m6A readers identified by mass spectrometry, control or m6A oligos (Synthesized by biomers.net GmbH, listed in Table S3) coupled to Streptavidin beads were incubated as above with either bacterial lysates expressing the respective recombinant proteins (GST- Gis2, Full length GST-Pho92 or truncation mutants of GST-Pho92) or SK1 yeast meiotic lysates expressing V5-tagged Gis2. After washes with protein incubation buffer, proteins bound to the beads were eluted by boiling for 5 minutes in laemmli buffer. The eluates were loaded onto SDS/ Polyacrylamide gels and assessed either by Coomassie staining or western blotting.

**m6A individual nucleotide resolution UV crosslinking and immunoprecipitation (miCLIP)**

The miCLIP was adapted from (Linder *et al*, 2015), while using the iiCLIP library preparation protocol (Lee *et al*, 2021). Total RNA extracted as above was then used for polyA selection with oligodT dynabeads (cat no 61005, Life Technologies) according to manufacturer's instructions and up to 10 µg polyA RNA was fragmented to around 150-200 nt with fragmentation reagent (AM8740) in a 20 µl reaction and heat to 70 °C for 4 min. The reaction is then stopped by addition of stop solution provided included in the kit. 4 ug mRNA was added to 450 µl immunoprecipitation buffer IP (50 Mm Tris pH 7.4, 100 mM NaCl, 0.05% NP-40) and incubated with 5 µg anti-m6A (ab151230) for 2h at 4 °C with rotation. polyA RNA was cross-linked twice to the antibody with 2x0.15 J/cm$^2$ UV light (254 nm) in a Stratalinker and then incubated with 30 µl protein G beads for 1.5 h at 4 °C with rotation. Bead bound antibody-RNA complexes recovered on a magnetic stand and washed twice with high salt buffer (50 mM Tris pH 7.4, 1 M NaCl, 1 mM EDTA, 1% NP-40, 0.1% SDS, 0.5% deoxycholate), twice with immunoprecipitation buffer and twice with polynucleotide kinase PNK buffer (20 mM Tris, 10 mM $MgCl_2$, 0.2 % Tween 20). Beads were resuspended in 20 µl of the following mixture: 4 µl 5x PNK pH 6.5 buffer (350 mM Tris-HCl, pH 6.5, 50 mM MgCl2, 5 mM dithiothreitol), 0.5 µl PNK (NEB M0201L), 0.5 µl RNasin, 15 µl water and incubated for 20 min at 37°C in thermomixer at 1100rpm. Then washes with 1 ml PNK buffer, 1 ml high-salt wash, 1 ml PNK buffer followed and beads were resuspended in 100 µl PNK buffer, moved to a new tube, place on magnet, removed supernatant and washed again once with 1 ml PNK buffer. Supernatant was removed and beads were resuspended in 25 µl of the ligation mix: 7.55 µl water, 3 µl 10X ligation buffer [500 mM Tris-HCl, pH 7.5, 100 mM MgCl2), 0.8 µl 100% DMSO, 2.5 µl RNA ligase –high concentration (M0437 NEB)], 0.5 µl PNK (NEB M0201L), 0.4 µl RNasin (NEB), 1.25 µl pre-adenylated adaptor L3-App (AGATCGGAAG_1,AGCGGTTCAG_2, 20 µM), 9 µl 50% PEG8000 and incubated overnight at 16°C in thermomixer at 1100rpm. Washes followed with

2x 1 ml high salt wash and 1 ml PNK buffers. PNK buffer was removed and 20 µl of the removal mix [12.5µl Nuclease-free H2O, 2µl NEB Buffer 2, 0.5ul Deµdenylase (NEB M0331S), 0.5µl RecJ endonuclease (NEB M0264S), 0.5µl RNasin, 4µl PEG400] added. Incubation followed for 1hr at 30°C, then for 30 min at 37°C whilst shaking at 1100rpm. The beads were washed with 2x1 ml high-salt wash and 1 ml PNK buffer. Then the sample was split in 2 fractions. 10% was labelled with [gamma-P32] ATP (($\gamma$-$^{32}$P)-ATP, SRP-301, Hartmann Analytic) in order to visualize mRNA-antibody complexes, the rest was not labelled and used for library preparation. 10% of the beads were collected, 4 µl of hot PNK mix [0.2 µl PNK (NEB M0201L), 0.5 µl 32P-γ-ATP, 0.4 µl 10x PNK buffer (NEB), 2.9 µl water] was added after removal of supernatant and incubated for 5min at 37°C in thermomixer at 1100rpm. Supernatant was removed and 100µl PNK buffer was added, incubated in thermomixer at 37°C /1100rpm for 1min, and again supernatant was removed.

SDS-PAGE and Western Blotting of labelled and non-labelled fractions followed with 4-12% NuPAGE Bis-Tris gel and MOPS buffer from Invitrogen and nitrocellulose membrane. The part of the membrane containing the signal was determined from the labelled fraction but the membrane was excised (75-150 kDa) from the non-labelled lane and cut into several pieces. 10µl proteinase K (Roche, 03115828001) was added in 200 µl PK+SDS (10 mM Tris-HCl, pH 7.4, 100 mM NaCl, 1 mM EDTA, 0.2% SDS) buffer and to the nitrocellulose piece and incubation followed with shaking at 1100 rpm for 60 min at 50°C. The solution was moved to another tube, 200 µl Phenol:Chloroform:Isoamyl Alcohol (Sigma P3803) added, mixed, and added to a pre-spun 2 ml Phase Lock Gel Heavy tube (713-2536, VWR). Incubation followed for 5 min at 30°C with shaking at 1100 rpm. The phases were separated by centrifugation for 5 min at 13000 rpm at room temperature. 800 µl chloroform was added to the top phase of the Phase Lock Gel Heavy tube and after centrifugation for 5 min at 13000 rpm at 4 °C the aqueous layer was transferred into a new tube. The RNA was precipitated by addition of 0.75 µl glycoblue (Ambion, 9510), 20 µl 5 M sodium chloride and 500µl 100% ethanol, followed by overnight incubation at -20°C and spin for 20 min at 15000 rpm at 4°C. The pellet was washed with 0.9 ml 80% ethanol and resuspended in 5.5 µl $H_2O$.

1 µl primer irCLIP_ddRT_## 1 pmol/µl and 0.5 µ l dNTP mix (10 mM) were added to the resuspended 5.5 µl RNA pellet. Incubation at 65°C for 5 min followed. Then the RT mix was added, 2 µl 5x SSIV buffer (Invitrogen), 0.5 µl 0.1 M DTT, 0.25 µl RNasin, 0.25 µl Superscript IV (Invitrogen), and the RT reaction followed (25°C 5 min, 50°C 5 min, 55°C 5 min). 1 µl of RnaseH (NEB, M0297S): RnaseA (Ambion, AM2274) mix was added and incubated at 37°C for 15 minutes. Agencourt AMPure XP beads (3x the volume of the RT reaction) added directly to the RT reaction and isopropanol to 1.7xvolume of the RT reaction. After 5 min incubation at RT, the beads were collected on magnetic stand, removed supernatant, 200µl 85% ethanol added and incubated for 30sec, then removed the supernatant. The last step was repeated and the beads were left to dry on the magnetic stand and finally eluted with 8 µl water.

Circularization followed with addition of 2 µl of circligase II mix (1 µl 10x CircLigase Buffer II , 0.5 µl CircLigase II, 0.5 µl 50mM MnCl2 – kit from Epicentre) and the reaction incubated at 60 $^{o}$C for 2 h. Ampure bead clean up was repeated as above and the cDNA eluted in 10µl water. The PCR reaction [1 µl cDNA, 0.25 µl primer mix

P5Solexa/P3Solexa 10 μM each, 5 μl Phusion HF Master mix (M0531S), 3.75 μl water] was performed for 20 cycles with the following programme 98°C 40 sec, 20x(98°C 20 sec/65°C 30 sec/72°C 45 sec), 72°C 3 min, the products run on 6% TBE gel and stained with SYBR green I.

cDNA was excised in the range of 145-400 nt and the gel fragment was placed inside a 1.5 ml tube with holes in the bottom and placed in a 2 ml collection tube. After centrifugation at 13000rpm for 5 min, 500μl of crush-soak gel buffer (500mM NaCl, 1mM EDTA, 0.05% SDS) was added to the gel and incubated at 65ºC for 2 h with thermomixer settings of 15 s at 1000 rpm, 45s rest. The liquid portion of the supernatant was transferred into a Costar SpinX column (Corning Incorporated, 8161) into which two 1 cm glass pre-filters (Whatman 1823010) were placed and centrifuged at 13000 rpm for 1 min. The solution was collected and 1 μl glycoblue, 50 μl 3M sodium acetate, pH 5.5 and 1.5 ml 100% ethanol were added. The RNA was precipitated overnight at -20ºC followed by centrifugation for 30 min at 15000 rpm at 4°C. The cDNA pellet was washed once with 0.9 ml 80% EtOH, resuspended in 10-15 μl RNase free $H_2O$ and submitted for sequencing.

The entire protocol was also performed with 1 μg input mRNA omitting IP, P32 labelling, SDS-PAGE/western blotting steps. We moved from adapter removal to reverse transcription. An RNA clean up step using RNA clean and concentrator kit (Zymo Research R1016) was included each time after adapter ligation and adapter removal.

**UV-C (254 nm) and 4-ThioUracil-UV-A (365 nm) individual nucleotide resolution UV Crosslinking and Immunoprecipitation (iCLIP)**

We used the improved iCLIP (iiCLIP) protocol (Lee *et al.*, 2021). 250 ml of SK1 cells were cultured as stated above and grown for 4h in SPO. For UV-A 4-TU-iCLIP, SK1 cells expressing the *FUR4* transgene (to facilitate uptake of 4-thiouracil) were grown for 4h in SPO containing 0.5 mM 4-thio uracil. After 4 hours, cells were collected in 10- or 20-ml PBS in a petri dish on ice and crosslinked with 0.8 J/cm$^2$ UV light (254 nm) or 12 J/cm$^2$ UV light (365 nm) in a Stratalinker 2400. Crosslinked cells were collected and resuspended in 2ml lysis buffer (50 mM Tris-HCl pH 7.5, 100 mM NaCl, 0.5% Na-DOC, 0.1% SDS, 0.5% NP-40, 1 mM DTT, 1X Protease inhibitor cocktail (HALT)) and pebbles prepared in liquid nitrogen. Cells were subjected to cryogenic lysis by freezer mill grinding under liquid nitrogen (SPEX 6875D Freezer/Mill, standard program: 15 cps for 6 cycles of 2 minutes grinding and 2 minutes cooling each). The resultant yeast “grindate” powder was resuspended in equal volume of lysis buffer and left to tumble in the cold for an hour. The cell lysate was initially clarified by a short 5-minute centrifugation at 1500 rpm at 4 degrees and then further purified by ultra-centrifugation at 45000 rpm for 1 hour at 4 degrees. The clarified lysates were subjected to DNase (2 µl of Turbo DNase per mg protein) and RNase (0.05 units/ mg lysate) treatment for 3 minutes at 37 degrees and then transferred to ice. Dynabeads (Protein G) were washed twice in lysis buffer before incubating 10 ug of V5 antibody (for each IP) with the beads for one hour at room temperature. Subsequently the beads were washed thrice in lysis before adding the lysates to the beads-antibody mix. The immunoprecipitation was carried out overnight at 4 degrees and from here onwards the same steps as in the miCLIP methods given above were followed starting with the wash with high salt buffer and including the library preparation with the only exception that RNA-protein complexes in this case were excised out of the membrane from above the size of the respective protein bands.

**iCLIP/miCLIP pre-processing**

Reads were demultiplexed using iCount demultiplex, and subsequently trimmed for adapter sequences and also for PHRED score > 30 using Trim Galore! (Krueger, 2017). Due to the high ncRNA content in miCLIP libraries a sequential mapping strategy was used for all libraries, which is available as a Snakemake (Koster & Rahmann, 2012) pipeline from www.github.com/ulelab/ncawareclip. Mapping was first to representative *Saccharomyces cerevisiae* snRNA and rRNA sequences downloaded from NCBI (Pruitt *et al*, 2005), followed by mature tRNA sequences (3′ CCA and 5’ G added) and the SacCer3 mitochondrial chromosome before being mapped to the SK1 MvO genome (available from http://cbio.mskcc.org/public/SK1_MvO/). PCR duplicates were removed using the unique molecular identifiers (UMIs). The start positions of uniquely mapping reads were taken as crosslinks. Detailed annotations were taken from (Chia *et al.*, 2021). tRNA annotations were downloaded from UCSC table browser, which sources the annotations from GtRNAdb (Chan & Lowe, 2016; Karolchik *et al*, 2004). SK1 annotations were supplemented with 3' and 5' UTR annotations derived (Chia *et al.*, 2021). All genome browser screenshots display stranded crosslinks per million (CPM) normalised bigWig files. Crosslinks at each position were divided by the total number of genomic crosslinks in the sample multiplied by one million.

**iCLIP/miCLIP Differential Analysis**

Peaks were called on iCLIP and miCLIP data with Clippy v1.2.0 (https://github.com/ulelab/clippy/releases/tag/v1.2.0) specifically with settings -n 20 -up 50 -down 100 -x 2.75 -hc 0.8 -mg 10 -mb 10 --no_exon_info. Reproducible, high quality replicate samples, as determined by principal component analysis (PCA), replicate correlations, library size and regional crosslink locations, were combined for peak calling and then individual sample coverage over these peaks was calculated using Bedtools map (Quinlan, 2014). Peaks were filtered to have at least 5 cDNAs in 3 WT or 3 *ime4*-∆ replicates, to come to a preliminary list of binding sites.

To determine Ime4p-dependence, WT vs. *ime4*∆ samples for Pho92 iCLIP, Gis2 iCLIP and m6A miCLIP were compared using DeSeq2, whilst controlling for gene expression changes by including measurements from mock miCLIP samples as a contrast in the linear model (Love *et al.*, 2014; McIntyre *et al*, 2020). Genes with less than 20 cDNA counts across 3 replicates were discarded from the analysis. P values were calculated using a likelihood-ratio test. A stringent threshold of log2FoldChange <= -2 and adjusted p value of < 0.001 was used to determine differentially bound sites for Pho92 and Gis2 iCLIP. For m6A miCLIP a threshold of log2FoldChange <= -1 and adjusted p value of < 0.001 were used. Due to the high depth of the iCLIP datasets, sites were further filtered based on a DeSeq2 base mean > 200, which is a measure of coverage in both iCLIP and mock iCLIP samples. The value is calculated as the average of the normalized cDNA counts per peak from all samples, divided by their size factors. Peak assignment to transcriptomic regions was performed using the following hierarchy to resolve any overlapping annotations: snoRNA > ncRNA >

STOP codon > 3' UTR > 5'UTR > last 100nt CDS > first 100nt CDS > CDS > intergenic.

**Published m6A sites definition**

To come to a group of published m6A sites to compare our data against, all Mazter-Seq sites with confidence group > 1 and all m6A-Seq sites were taken (Garcia-Campos *et al*, 2019; Schwartz *et al*, 2013). To robustly map these to the MvO SK1 genome assembly, all intervals were expanded to 150nt, sequences retrieved and BLAST used to get mappings. Mappings were filtered to those that were unique and in the case of Mazter-Seq, perfectly aligned to an "ACA" sequence. To create a consensus list, any m6A-seq region that overlapped with Mazter-Seq site(s) was removed, under the assumption that the signal was representing the sites detected at higher resolution by Mazter-Seq - although it is possible that adjacent non-ACA sequence context m6A sites would not be represented in the list. This procedure resulted in a list of 1297 m6A regions.

**Distance to nearest m6A**

Distances between peak sets were calculated using bedtools closest, with parameters -s -t first -d (Quinlan, 2014).

**Go term enrichment**

Gene list enrichment analysis was performed using YeastEnrichr, specifically using KEGG 2019 pathways (https://maayanlab.cloud/YeastEnrichr/) (Chen *et al*, 2013; Kuleshov *et al*, 2016) .

### Motif analysis

Motifs were discovered in peak regions by resizing all peaks to 100nt and obtaining their fasta sequences to submit to STREME (Bailey, 2021). Either shuffled sequences or another peak set were used as background, as indicated in the main text. Motifs were plotted around the centre of peaks using a custom script available upon request.

### Metaprofiles

Bigwigs were generated from bedgraphs using UCSC BedGraphToBigwig (Kuhn *et al*, 2013) and metaprofiles were generated around regions of interest using DeepTools ComputeMatrix and PlotHeatmap (Ramirez *et al*, 2014). Further integration and plotting was performed using R with ggplot2, dplyr, data.table, stringr and cowplot packages (Dowle, 2019; Wickham, 2010; Wickham, 2015, 2016; Wilke, 2019).

### Determining Pho92 intronic overlaps

Due to the sparsity of SK1 genome annotations, many known introns that are present in SacCer3 annotations are not present in SK1 annotation. Therefore, to determine intronic overlaps, all Pho92-bound genes with introns in SacCer3 annotations were manually inspected alongside miCLIP-input and checked for whether Pho92 peaks overlapped regions suspected to be introns due to: a) sharp drop in miCLIP-input signal in the region, b) corresponding intron in SacCer3 annotations.

### Immunoprecipitation and mass spectrometry

Extracts were prepared in a mild buffer (50 mM Tris-HCl pH 7.5, 100 mM NaCl, 10 mM MgCl2, 0.1% Na-DOC, 0.1% SDS, 0.1% NP-40, 1 mM DTT, 1X Protease inhibitor cocktail (HALT)) using cryogenic lysis as detailed above from S288C expressing *pCYC1*- FLAG-Pho92, *pPYK1*- FLAG-Pho92 and SK1 expressing Pho92-V5. Untagged S288C and SK1 strains were used as controls. Extracts were incubated with either Flag (M2-agarose) or V5 agarose for 2 hours at 4 degrees. Washes were done with the same buffer as above but with an addition of 0.2% NP-40 instead. Elutions from Flag beads were done with 2 mg/ml flag -peptide. All eluates were combined and samples were further processed using the PreOmics iST (inStageTip) protocol and the manufacturer's protocol was followed. Eluted peptides were dried by vacuum centrifugation. Samples were solubilised in 20 µl of 0.1 % TFA & injected in technical triplicate Orbitrap Fusion Lumos Tribrid mass spectrometer.

For V5 pull downs, after the V5 peptide elutions (which were found to be inefficient), a further laemmli buffer elution from the beads by boiling was performed. Laemmli buffer eluate samples were processed using the single-pot, solid phase-enhanced, sample preparation (SP3) technology. Paramagnetic beads were added to the tryptic digests and peptide eluates were purified using C18 stagetips. Samples were injected in technical triplicates on the QExactive instrument.

All raw files were analysed in MaxQuant v1.6.0.13 against the 2019 SwissProt *S. cerevisiae* protein database & additional data analyses were performed in Perseus v1.4.0.2.

**Co-Immunoprecipitation and GST pull downs**

Cells were resuspended in lysis buffer (50 mM Tris-HCl pH 7.5, 150 mM NaCl, 10 mM $MgCl_2$, 0.5% Na-DOC, 0.1% SDS, 0.2% NP-40, 1 mM DTT, and 1X HALT protease inhibitor) and lysed using Zirconia beads in beadbeater (BioSpec). The lysates were cleared via centrifugation (16,000g at 4°C for 20min). Flag-Pho92 and associated proteins were pulled down using anti-Flag M2-conjugated agarose beads by incubating at 4°C for 2 hours. Unbound proteins were washed from the beads with wash buffer (50 mM Tris-HCl pH 7.5, 1 M NaCl, 10 mM $MgCl_2$, 0.5% Na-DOC, 0.1% SDS and 0.5% NP-40). Bound proteins were eluted by competition with an excess of free Flag peptide. Samples were analysed by Western Blotting.

Soluble GST-tagged fusion proteins expressed and prepared as above were immobilized on glutathione-agarose beads and subsequently used for binding proteins from yeast extracts expressing HA-tagged Paf1C or Rap1 prepared as above for co-immunoprecipitations. The bound proteins were eluted off the GA beads and analyzed by Western blotting by probing with anti- HA antibody.

**Global whole-cell proteome profiling**

Cells were collected by centrifugation, re-suspended in cold 5% v/v TCA and incubated on ice for 30 min. Samples were washed with acetone, then completely air-dried. The dried pellet was resuspended in protein breakage buffer (50 mM Tris (pH 7.5), 1 mM EDTA, 2.75 mM dithiothreitol (DTT), HALT protease inhibitors) and disrupted using 0.5 mm glass beads for 2 minutes in a Mini Beadbeater (Biospec). To the lysate, 3X concentrated modified laemmli buffer (187.5 mM Tris (pH 6.8), 6.0% v/v β-mercaptoethanol, 30% v/v glycerol, 9.0% v/v SDS) was added along with protease inhibitors (HALT) to result in a final concentration of 62.5 mM Tris, 2% β-mercaptoethanol, 10% glycerol and 3% SDS and denatured at 95°C for 5 min.

Reduction and alkylation by the addition of 20mM TCEP and 40 mM chloroacetamide to the provided protein lysates was carried out for 10 min at 70°C. Protein amounts were confirmed, following an SDS–PAGE gel of 4% of each sample against an in-house cell lysate of known quantity. A volume corresponding to 50 μg of each sample was taken along for digestion. Proteins were precipitated overnight at −20°C after addition of a 4× volume of ice-cold acetone. The following day, the samples were centrifuged at 20,800 x g for 30 min at 4°C and the supernatant carefully removed. Pellets were washed twice with 1 ml ice-cold 80% (v/v) acetone in water then centrifuged at 20,800 x g at 4°C. They were then allowed to air-dry before addition of 120 μl of digestion buffer (1 M Guanadine Hydrochloride, 100 mM HEPES, pH8). Samples were resuspended with sonication, LysC (Wako) was added at 1:100 (w/w) enzyme:protein, and digestion proceeded for 4 h at 37°C with shaking (Eppendorf ThermoMixer®C, thermoblock for 1.5 ml tubes, at 1,000 rpm for 1 h, then 650 rpm). Samples were then diluted 1:1 with Milli-Q water, and trypsin (Pierce) added at the same enzyme to protein ratio. Samples were further digested overnight at 37°C with shaking (650 rpm). The following day, digests were acidified by the addition of TFA to a final concentration of 2% (v/v) and then desalted with Waters Oasis® HLB μElution Plate 30 μm (Waters Corporation, Milford, MA, USA) in the presence of a slow vacuum. In this process, the columns were conditioned with 3 × 100 μl solvent B (80% (v/v) acetonitrile; 0.05% (v/v) formic acid) and equilibrated with 3 × 100 μl solvent A (0.05% (v/v) formic acid in Milli-Q water). The samples were loaded, washed 3 times with 100 μl solvent A, and then eluted into with 50 μl solvent B. The eluates were dried down with the speed vacuum centrifuge and dissolved at a concentration of 1 μg/μl in reconstitution buffer (5% (v/v) acetonitrile, 0.1% (v/v) formic acid in Milli-Q water).

Digested peptides were separated using the Dionex U3000 nano UHPLC system (Thermo) fitted with a trapping (PepMap Acclaim C18, 5 μm, 0.2 mm x 20 mm) and an analytical column (EasySpray PepMap C18, 2 μm, 75 μm x 500 mm), both held at 40 ˚C. The outlet of the analytical column was coupled directly to an Orbitrap Fusion Lumos Tribrid mass spectrometer (Thermo Fisher Scientific) using the EasySpray source. Peptides were eluted via a non-linear gradient from 100% aqueous/0.1% Formic acid to 40% Acetonitrile/0.1% Formic Acid in 98 min. Total runtime was 120 min, including clean-up and column re-equilibration. The RF lens was set to 30%. The spray voltage was set to 2.2 kV and source temperature 275 ˚C. Default charge state was set to 4+. For data acquisition and processing Tune version 3.3 was employed. Data Independent Acquisition (DIA) data was acquired with full scan MS spectra over the mass range 350-1650 m/z in profile mode in the Orbitrap with resolution of 120,000 FWHM. The filling time was set at maximum of 20 ms with limitation of 1E6 ions. DIA MS2 scans were acquired with 34 mass window segments of differing widths across the MS1 mass range with a cycle time of 3 seconds. HCD fragmentation (30% collision energy) was applied and MS/MS spectra were acquired in the Orbitrap at a resolution of 30,000 FWHM over the mass range 200-2000 m/z after accumulation of 1E6 ions or after filling time of 70 ms (whichever occurred first. Data were acquired in profile mode.

DIA data were searched directly against the SwissProt Saccharomyces Cerevisiae database (6721 entries) and a list of common contaminants using Direct DIA in Spectronaut (version 14, Biognosys AG, Schlieren, Switzerland). The following modifications were included in the search: Carbamidomethyl (C) (Fixed) and Oxidation (M)/Acetyl (Protein N-term; Variable). A maximum of 2 missed cleavages for trypsin were allowed. The identifications were filtered to satisfy FDR of 1% on peptide and protein level. The DirectDIA analysis resulted in 33575 precursors and 3292 Protein Groups as full profiles across the 6 samples. Precursor matching, protein inference, and quantification were performed in Spectronaut using default settings. The candidate table was exported for further data visualisation.

**Chromatin immunoprecipitation**

Chromatin immunoprecipitation (ChIP) experiments were performed as described previously (Tam & van Werven, 2020). Cells were fixed in 1.0% v/v formaldehyde for 20 min at room temperature and quenched with 100 mM glycine. Cells were lysed in FA lysis buffer (50 mM HEPES–KOH, pH 7.5, 150 mM NaCl, 1mM EDTA, 1% Triton X-100, 0.1% Na-deoxycholate, 0.1% SDS and protease cocktail inhibitor (complete mini EDTA-free, Roche)) using beadbeater (BioSpec) and chromatin was sheared by sonication using a Bioruptor (Diagenode, 9 cycles of 30 s on/off). Extracts were incubated for 2 h at room temperature with anti-V5 agarose beads (Sigma), washed twice with FA lysis buffer, twice with wash buffer 1 (FA lysis buffer containing 0.5M NaCl), and twice with wash buffer 2 (10 mM Tris–HCl, pH 8.0, 0.25M LiCl, 1 mM EDTA, 0.5% NP-40, 0.5% Na-deoxycholate). Subsequently, reverse cross-linking was done in 1% SDS-TE buffer (100 mM Tris pH 8.0, 10 mM EDTA, 1.0% v/v SDS) at 65°C overnight. After 2 h of proteinase K treatment, samples were purified, and DNA fragments were quantified by real-time PCR using PowerUP SYBR green master mix (Thermo Fisher Scientific) using primers described in Table S3.

**Fluorescence microscopy and quantification**

Cells were collected from sporulation cultures at desired time points, pelleted via centrifugation (1500g, 2min) and resuspended in approximately 100 µl of SPO media. Live cell image acquisition was conducted using a Nikon Eclipse Ti inverted microscope. Exposure times were set as follows: 500ms brightfield, 50ms GFP, and 50ms mCherry. An ORCA-FLASH 4.0 camera (Hamamatsu) and NIS-Elements AR software (Nikon) were used to collect images. Quantification of fluorescence signals was performed using FiJI software (Schindelin *et al*, 2012). ROIs were manually drawn around the periphery of each cell. The mean intensity in each channel per cell was multiplied by the cell area to obtain mean signal. The signal for each channel was corrected for cell-free background fluorescence in a similar way. Values for nuclear protein localisation were derived via the division of nuclear / whole cell signal. Whole cell value ranges were set for each time point between different strains for a fair comparison of nuclear signals. For the analyses, 150 cells were quantified per sample.

**Ribosome association by sucrose cushion**

Ribosomal fractions were separated from soluble components by centrifugation through sucrose cushions (Trotter *et al*, 2008). Cell extracts were prepared from S288C 100 ml cells $OD_{600}$ 0.5 in YPD, collected and centrifuged after addition of 10 µg/ml cycloheximide. Cell pellets were washed with 1 ml CSB buffer [300 mM sorbitol, 20 mM Hepes (pH 7.5), 1 mM EGTA, 5 mM $MgCl_2$, 10 mM KCl, 10% (v/v) glycerol and 10 µg/ml cycloheximide, protease inhibitors] and resuspended in 500 µl

CSB buffer, glass beads added and breakage followed in a bead beater for 3 min. Additional 300 µl CSB buffer were added and cell suspension was centrifuged for 4 min at full speed and 4ºC. The supernatant was moved to a fresh tube and centrifuged at 10000 rpm for 15 min at 4ºC. 500 µl of supernatant was layered onto 400 µl 60 % sucrose in CSB without sorbitol and centrifuged at 5500 rpm for 3.5 h in TLA110 at 4ºC. The supernatant was precipitated with 20% TCA and this as well as the ribosomal pellet were resuspended each in 50 µl Laemmli sample buffer.

**Ribosome fractionation on sucrose gradients**

Performed as previously described (Ashe *et al*, 2000). SK1 Cells were grown till saturation for 24h in YPD, then diluted at $OD_{600}$ = 0.4 to pre-sporulation medium BYTA grown for about 16h, subsequently centrifuged, washed with sterile Milli-Q water, centrifuged again, re-suspended at $OD_{600}$ = 1.8 in sporulation medium (SPO) and incubated at 30°C. Yeast cells were harvested after 4h in SPO. S288C cells were collected at $OD_{600}$ 0.5 in YPD as for sucrose cushions. 200ml cells were collected with addition of cycloheximide to 100 µg/ml final concentration, washed and resuspended in 2 ml lysis buffer (20 mM Hepes pH 7.4, 2 mM MgAcetate, 100 mM KAcetate, 100 µg/ml cycloheximide, 500 µM DTT, PIC, RNase inhibitor), and pebbles prepared in liquid nitrogen. Cells were subjected to cryogenic lysis by freezer mill grinding under liquid nitrogen (SPEX 6875D Freezer/Mill, standard program: 15 cps for 6 cycles of 2 minutes grinding and 2 minutes cooling each). Yeast “grindate” powder was stored at -80 °C. 0.5 ml yeast grindate powder was resuspended in 0.5 ml lysis buffer and centrifuged at 10000 rpm for 5 min at 4°C. Supernatant moved to a fresh tube and centrifuged at 10000 rpm for 15 min at 4°C. 300 ug of $A_{260}$ was loaded onto a 10-45% sucrose gradient. The sucrose solutions were in the following buffer: 10 mM Tris Acetate pH 7.4, 70 mM ammonium acetate, 4 mM MgAcetate. Polysome profiles were obtained from a Biocomp fractionator and samples collected for further analysis. For protein analysis 20 % TCA was added to the fractions collected, protein precipitated and run on western blotting. For RNA-seq, polysome fractions were precipitated overnight at -20°C with 400 µl isopropanol. The RNA pellet was precipitated and resuspended in 50 µl RNase free water, 800 µl TES and 800 µl Acid phenol (see RNA extraction-miclip section) added and incubated for 5 min at RT. Further the RNA pellet was precipitated, DNase treated and column purified as in RNA extraction/miclip section.

**Next generation sequencing**

The NEBNext® Ultra™ II Directional RNA Library Prep Kit for Illumina with the NEBNext Poly(A) mRNA Magnetic Isolation Module was used. Approximately 12 ng of RNA was used as input, according to manufacturer's instructions. Sequencing was carried out on the Illumina Hiseq 4000 or NovaSeq 6000 with single ended 100 bp reads.

**m6A-mRNA quantification by LC-MS/MS and ELISA**

RNA extracted and DNase treated as described above and polyA RNA isolated after purification twice with oligodT dynabeads (Ambion, 61005) according to manufacturer's instructions. 375 ng polyA selected RNA in 20.5 µl H2O was digested with addition of 2.5µl of 10X Buffer (25mM ZnCl2, 250mM NaCl, 100mM NaAcetate), 2µl of Nuclease P1 (1U/µl) (Sigma, N8630) and incubated for 4 h at 37 °C. Then 3µl of 1M ammonium bicarbonate, 1µl Alkaline Phosphatase (NEB, M0525) and 1µl of

H2O were added. Incubation followed for 2 h at 37 °C. 19 µl H2O and 1 µl of 5 % Formic Acid added to the reaction mixture and filtered with a 1.5ml microfuge tube containing a 0.22µM inside filter. The sample was then injected (20 µl) and analysed by LC-MS/MS using a reverse phase liquid chromatography C18 column and a triple quadrupole mass analyser (Agilent 6470 or Thermo Scientific TSQ Quantiva) instrument in positive electrospray ionisation mode. Flow rate was at 0.2 ml/min and column temperature 25 °C with the following gradient: 2 min 98% eluent A (0.1% formic acid and 10 mM ammonium formate in water) and 2% eluent B (0.1% formic acid and 10 mM ammonium formate in MeOH), 75% A and 25% B up to 10 min, 20% A and 80% B up to 15 min, 98% A and 2% B up to 22.5 min. Nucleosides were quantified using the mass transitions of 281 to 150 for $m^6$A and 268 to 136 for adenosine and a calibration curve of pure nucleosides standards.

For ELISA we either used the EpiQuik m6A RNA methylation quantification kit from EpiGentek (P-9005) according to manufacturer's protocol or the protocol we developed. In our method 90 µl binding solution (ab156917) and 50 ng of twice polyA selected RNA (similar to LC-MS/MS above) were added to each well of a 96 well plate (ab210903). Three technical replicates were used for each sample and standard. The plate incubated at 37 °C for 2h. After incubation, binding solution and mRNA removed and wells washed 4 times with 200 µl PBSTween (0.1%) in each wash. 100 µl of primary antibody (mRabbit A19841, 1:10000 in PBSTween) containing 0.5 µg/ml total RNA from *ime4*Δ mutant added and the plate incubated at RT for 1 h. Following incubation, solution was removed and wells washed 4 times with 200 µl PBSTween (0.1%) each time, while incubating the plate with the third wash for 5 min at RT. 100 µl of secondary antibody (anti-rabbit 1:5000 in PBST, ab205718) added and incubation at RT for 30 min followed. Solution was removed and wells washed 5 times with 200 µl PBSTween (0.1%) each time, while incubating the plate with the third wash for 5 min at RT. 100 µl of developing solution (ab156917) added to the wells and left for 20-30 min before addition of 100 µl of stop solution (ab171524). Absorbance was recorded at 450 nm. Quantity of m6A was calculated using a calibration curve with standards of in vitro transcribed m6A-RNA and adenosine-RNA in the range of 0.0005-0.0125 ng for m6A-RNA with addition of 50 ng adenosine-RNA in each standard. All reagents bought from Abcam. In vitro transcribed adenosine-RNA and m6A-RNA were synthesized with MEGAscript T7 Transcription kit (AM1333, Ambion) from ATP and $N^6$-Methyl-ATP (NU1101L, 2B Scientific) respectively.

**m6A-mRNA stability assay**

SK1 cells were grown till saturation for 24h in YPD, then diluted at OD600 = 0.4 to pre-sporulation medium BYTA grown for about 16 h, subsequently centrifuged, washed with sterile Milli-Q water, centrifuged again, re-suspended at OD600 = 1.8 in SPO. After 4 h in SPO cultures were treated either with 3 µg/ml thiolutin (T2834, Cambridge Bioscience) or with 3 µg/ml thiolutin and 100 µg/ml cycloheximide (C4859, Merck) and samples collected at indicated timepoints. RNA extraction, polyA selection and m6A quantification either with ELISA or LC-MS/MS followed as described in relevant sections.

**Statistical analyses**

Data statistics and statistical analyses indicated in the figure legends were computed using GraphPad Prism version 8.2.0 for Windows, GraphPad Software, San Diego, CA, USA, www.graphpad.com. Data from the imaging experiments were analysed using paired parametric two-tailed Welch's t test with 95% confidence. Boxplots highlight median and quartiles 1 and 3. p Values are indicated in the figures, where ns stands for no significant difference, **p<0.01.

**Data availability**

The miCLIP, iCLIP and RNA-seq RAW and processed data are available to review GEO accession GSE193561:

Go to https://eur03.safelinks.protection.outlook.com/?url=https%3A%2F%2Fwww.ncbi.nlm.nih.gov%2Fgeo%2Fquery%2Facc.cgi%3Facc%3DGSE193561&data=04%7C01%7C%7Cd763bb9f0b064c29195c08d9d9935295%7C4eed7807ebad415aa7a99170947f4eae%7C0%7C0%7C637780049042619595%7CUnknown%7CTWFpbGZsb3d8eyJWIjoiMC4wLjAwMDAiLCJQIjoiV2luMzIiLCJBTiI6Ik1haWwiLCJXVCI6Mn0%3D%7C3000&sdata=1DrdEauEN%2B50wKJrzoaDWAS4JiJEWzoCiQJM3MdYVTg%3D&reserved=0

Enter token anmtmqgotlsfrsl into the box
