## Supplementary material for "m6A reader Pho92 is recruited co-transcriptionally and couples translation efficacy to mRNA decay to promote meiotic fitness in yeast": Table S2

**Table S2 . Plasmids used in this study**

| Plasmid No. | Name |
| --- | --- |
| FW P768 | pGEX-6P-1 |
| FW P769 | pGEX-6P-1-Gis2 |
| FW P759 | pGEX-6P-1-Pho92-FL |
| FW P760 | pGEX-6P-1-Pho92-NTDd |
| FW P761 | pGEX-6P-1-Pho92-YTHd |
| FW P718 | pK3FS-PYK1 (Addgene Plasmid #85777) |
| FW P719 | pK3FS-CYC1 (Addgene Plasmid #85779) |
